# Inferring the timing and strength of natural selection and gene migration in the evolution of chicken from ancient DNA data

**DOI:** 10.1101/2021.04.30.442150

**Authors:** Wenyang Lyu, Xiaoyang Dai, Mark Beaumont, Feng Yu, Zhangyi He

## Abstract

With the rapid growth of the number of sequenced ancient genomes, there has been increasing interest in using this new information to study past and present adaptation. Such an additional temporal component has the promise of providing improved power for the estimation of natural selection. Over the last decade, statistical approaches for detection and quantification of natural selection from ancient DNA (aDNA) data have been developed. However, most of the existing methods do not allow us to estimate the timing of natural selection along with its strength, which is key to understanding the evolution and persistence of organismal diversity. Additionally, most methods ignore the fact that natural populations are almost always structured, which can result in overestimation of the effect of natural selection. To address these issues, we introduce a novel Bayesian framework for the inference of natural selection and gene migration from aDNA data with Markov chain Monte Carlo techniques, co-estimating both timing and strength of natural selection and gene migration. Such an advance enables us to infer drivers of natural selection and gene migration by correlating genetic evolution with potential causes such as the changes in the ecological context in which an organism has evolved. The performance of our procedure is evaluated through extensive simulations, with its utility shown with an application to ancient chicken samples.

## 1. Introduction

With modern advances in ancient DNA (aDNA) techniques, there has been a rapid increase in the availability of time serial samples of segregating alleles across one or more related populations. The temporal aspect of such samples reflects the combined evolutionary forces acting within and among populations such as genetic drift, natural selection and gene migration, which can contribute to our understanding of how these evolutionary forces shape the patterns observed in contemporaneous samples. One of the most powerful applications of such genetic time series is to study the action of natural selection since the expected changes in allele frequencies over time are closely related to the timing and strength of natural selection.

Over the past fifteen years, there has been a growing literature on the statistical inference of natural selection from time series data of allele frequencies, especially in aDNA (see Malaspinas, 2016; Dehasque et al., 2020, for excellent reviews). Typically, estimating natural selection from genetic time series is built on the hidden Markov model (HMM) framework proposed by Bollback et al. (2008), where the allele frequency trajectory of the underlying population through time was modelled as a latent variable following the Wright-Fisher model introduced by Fisher (1922) and Wright (1931), and the allele frequency of the sample drawn from the underlying population at each sampling time point was modelled as a noisy observation of the latent population allele frequency. In their likelihood computation, the Wright-Fisher model was approximated through its standard diffusion limit, known as the Wright-Fisher diffusion, which was then discretised for numerical integration with a finite difference scheme. Their approach was applied to analyse the aDNA data associated with horse coat colouration in Ludwig et al. (2009) and extended to more complex evolutionary scenarios (see, *e*.*g*., Malaspinas et al., 2012; Steinrücken et al., 2014;Ferrer-Admetlla et al., 2016; Schraiber et al., 2016; He et al., 2020b,c).

Natural populations are almost always structured, which affects the relative effect of natural selection and genetic drift on the changes in allele frequencies over time. This can cause overestimation of the selection coefficient (Mathieson et al., 2015). However, all existing methods based on the Wright-Fisher model for the inference of natural selection from temporally-spaced allele frequency data lack the ability to account for the confounding effect of gene migration, with the exception of Mathieson & McVean (2013), which could model population structure. Mathieson & McVean (2013) is an extension of Bollback et al. (2008) for the inference of metapopulations,which enables the co-estimation of the selection coefficient and the migration rate from genetic time series. However, their method could become computationally cumbersome for large population sizes and evolutionary timescales since their likelihood computation was carried out with the Wright-Fisher model. High computational costs for large evolutionary timescales become a strong limitation for the analysis of aDNA.

More recently, Loog et al. (2017) introduced a Bayesian statistical framework for estimating the timing and strength of selection from genetic time series while explicitly modelling migration from external sources. Their approach also allows the co-estimation of the underlying population allele frequency trajectory through time, which is important for understanding the drivers of selection. However, the population size is assumed to be large enough in their method to ignore genetic drift, which simplifies the likelihood computation but restricts the application to aDNA. In this work, we develop a novel HMM-based approach for the Bayesian inference of natural selection and gene migration to re-analyse the ancient chicken samples from Loog et al. (2017). Our method is built upon the HMM framework of Bollback et al. (2008), but unlike most existing approaches, it allows the joint estimation of the timing and strength of selection and migration. Such an advance enables us to infer the drivers of selection and migration by correlating genetic evolution with ecological and cultural shifts. Our main innovation is to introduce a multi-allele Wright-Fisher diffusion for a single locus evolving under natural selection and gene migration, including the timing of selection and migration. This diffusion process characterises the allele frequency trajectories of the underlying population over time, where the alleles that migrate from external sources are distinguished from those that originate in the underlying population. Such a setup allows a full using of available quantities such as the proportion of the modern European chicken that have Asian origin as a direct result of gene migration. Our posterior computation is carried out through the particle marginal Metropolis-Hastings (PMMH) algorithm of Andrieu et al. (2010) with blockwise sampling, which permits a joint update of the underlying population allele frequency trajectories. We evaluate the performance of our procedure through extensive simulations, with its utility shown with an application to the ancient chicken samples.

## 2. Materials and Methods

In this section, we first introduce the multi-allele Wright-Fisher diffusion for a single locus evolving under natural selection and gene migration and then present our Bayesian method for the joint inference of selection and migration from time series allele frequency data.

### 2.1. Wright-Fisher diffusion

Let us consider a population of *N* randomly mating diploid individuals at a single locus 𝒜 with discrete and nonoverlapping generations, where the population size *N* is finite and fixed over time. Suppose that at locus 𝒜 there are two allele types, labelled 𝒜_1_ and 𝒜_2_, respectively. We attach the symbol 𝒜_1_ to the mutant allele, which arises only once in the population and is positively selected once the evolution starts to act through selection, and we attach the symbol 𝒜_2_ to the ancestral allele, which originally exists in the population.

According to Loog et al. (2017), we characterise the population structure with the continent-island model (see, *e.g*., Hamilton, 2011, for an introduction). More specifically, the population is subdivided into two demes, the continental population and the island population. To distinguish between the alleles found on the island but emigrated from the continent or were originally on the island, the mutant and ancestral alleles that originated on the island are labelled 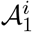 and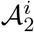, respectively, and the mutant and ancestral alleles that were results of emigration from the continent are labelled 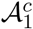 and 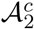, respectively. Such a setup enables us to trace the alleles that migrate from external sources evolving in the island population, thereby allowing the integration of available information such as the proportion of the modern European chicken that have Asian origin in the inference of selection. Suppose that the continent population is large enough such that gene migration between the continent and the island does not affect the genetic composition of the continent population. Our interests focus on the island population dynamics so in what follows the population refers to the island population unless noted otherwise.

To investigate the island population dynamics under natural selection and gene migration, we specify the life cycle of the island population, which starts with zygotes that selection acts on. Selection takes the form of viability selection, and the relative viabilities of all genotypes are shown in Table 1, where *s* ∈ [0, 1] is the selection coefficient, and *h* ∈ [0, 1] is the dominance parameter. After selection, a fraction *m* of the adults on the continent migrate into the population of mating adults on the island, which causes the change of the genetic composition of the island population, *i.e*., fraction *m* of the adults on the island are immigrants from the continent,and fraction 1 − *m* of adults were originally already on the island. The Wright-Fisher reproduction introduced by Fisher (1922) and Wright (1931) finally completes the life cycle, which corresponds to randomly sampling 2*N* gametes with replacement from an effectively infinite gamete pool to form new zygotes in the next generation through random union of gametes.

**Table 1:** Relative viabilities of all possible genotypes at locus 𝒜 when we distinguish between the alleles that originate on the island and the alleles that emigrate from the continent.

| | $\mathcal{A}_1^i$ | $\mathcal{A}_2^i$ | $\mathcal{A}_1^c$ | $\mathcal{A}_2^c$ |
| --- | --- | --- | --- | --- |
| $\mathcal{A}_1^i$ | 1 | $1 - hs$ | 1 | $1 - hs$ |
| $\mathcal{A}_2^i$ | $1 - hs$ | $1 - s$ | $1 - hs$ | $1 - s$ |
| $\mathcal{A}_1^c$ | 1 | $1 - hs$ | 1 | $1 - hs$ |
| $\mathcal{A}_2^c$ | $1 - hs$ | $1 - s$ | $1 - hs$ | $1 - s$ |

We let 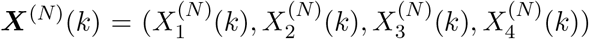 denote the frequencies of the 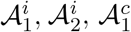 and 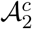alleles in *N* zygotes of generation *k* ∈ ℕ on the island, which follows the multi-allele Wright-Fisher model with selection and migration described in Supplemental Information, File S1. We assume that the selection coefficient and the migration rate are both of order 1*/*(2*N*) and fixed from the time of the onset of selection and migration up to present. We run time at rate 2*N, i.e*., *t* = *k/*(2*N*), and like Cherry & Wakeley (2003), we scale the selection coefficient and the migration rate as

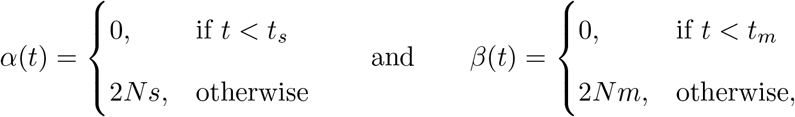

where *t*_*s*_ and *t*_*m*_ are the starting times of selection and migration on the island measured in the units of 2*N* generations. As the population size *N* tends to infinity, the Wright-Fisher model ***X***^(*N*)^ converges to a diffusion process, denoted by ***X*** = {***X***(*t*), *t* ≥ *t*_0_}, evolving in the state space 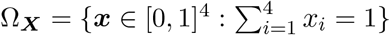 and satisfying the stochastic differential equation (SDE) of the form

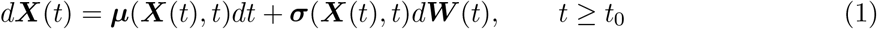

with initial condition ***X***(*t*_0_) = ***x***_0_. In Eq. (1), ***µ***(***x***, *t*) is the drift term

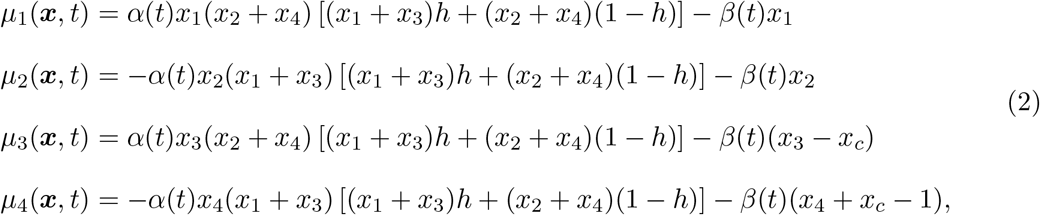

where *x*_*c*_ is the frequency of the 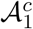 allele in the continent population, which is fixed over time, ***σ***(***x***, *t*) is the diffusion term

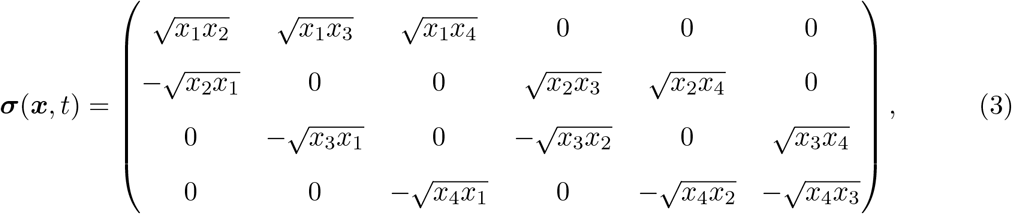

and ***W*** (*t*) is a six-dimensional standard Brownian motion. The explicit formula for the diffusion term ***σ***(***x***, *t*) in Eq. (3) is obtained by following He et al. (2020a). The proof of the convergence follows in the similar manner to that employed for the neutral two-locus case in Durrett (2008, p. 323). We refer to the stochastic process ***X*** = {***X***(*t*), *t* ≥ *t*_0_} that solves the SDE in Eq. (1) as the multi-allele Wright-Fisher diffusion with selection and migration.

### 2.2. Bayesian inference of natural selection and gene migration

Suppose that the available data are sampled from the underlying island population at time points *t*_1_ < *t*_2_ < … < *t*_*K*_, which are measured in units of 2*N* generations to be consistent with the Wright-Fisher diffusion timescale. At the sampling time point *t*_*k*_, there are *c*_*k*_ mutant alleles (*i.e*., the 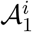 and 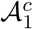 alleles) and *d*_*k*_ continent alleles alleles (*i.e*., the 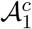 and 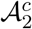 alleles) observed in the sample of *n*_*k*_ chromosomes drawn from the underlying island population. Note that in real data, the continent allele count of the sample may not be available at each sampling time point, *e.g*., the proportion of the European chicken that have Asian origin is only available in the modern sample (Loog et al., 2017). The population genetic parameters of interest are the scaled selection coefficient *α* = 2*Ns*, the dominance parameter *h*, the selection time *t*_*s*_, the scaled migration rate *β* = 2*Nm*, and the migration time *t*_*m*_, as well as the underlying allele frequency trajectories of the island population. For simplicity, in the sequel we let ***ϑ***_*s*_ = (*α, h, t*_*s*_) be the selection-related parameters and ***ϑ***_*m*_ = (*β, t*_*m*_) be the migration-related parameters, respectively.

#### 2.2.1. Hidden Markov model

We apply an HMM framework similar to the one proposed in Bollback et al. (2008), where the underlying population evolves under the Wright-Fisher diffusion in Eq. (1) and the observations are modelled through independent sampling from the underlying population at each given time point. Unlike Loog et al. (2017), we jointly estimate the timing and strength of selection and migration, including the allele frequency trajectories of the underlying population. Our Wright-Fisher diffusion can directly trace the temporal changes in the frequencies of the alleles in the island population that results from emigrants from the continent population. We can therefore make the most of other available information like the proportion of the modern European chicken with Asian ancestry in the most recent sample reported in Loog et al. (2017), which provides valuable information regarding the timing and strength of migration.

Let ***x***_1:*K*_ = (***x***_1_, ***x***_2_, …, ***x***_*K*_) be the allele frequency trajectories of the underlying population at the sampling time points ***t***_1:*K*_. Under our HMM framework, the joint posterior probability distribution for the population genetic quantities of interest and the allele frequency trajectories of the underlying population is

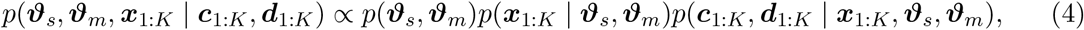

where *p*(***ϑ***_*s*_, ***ϑ***_*m*_) is the prior probability distribution for the population genetic quantities and can be taken to be a uniform distribution over the parameter space if their prior knowledge is poor, *p*(***x***_1:*K*_ | ***ϑ***_*s*_, ***ϑ***_*m*_) is the probability distribution for the allele frequency trajectories of the underlying population at the sampling time points ***t***_1:*K*_, and *p*(***c***_1:*K*_, ***d***_1:*K*_ | ***x***_1:*K*_, ***ϑ***_*s*_, ***ϑ***_*m*_) is the probability of the observations at the sampling time points ***t***_1:*K*_ conditional on the underlying population allele frequency trajectories.

With the Markov property of the Wright-Fisher diffusion, we have

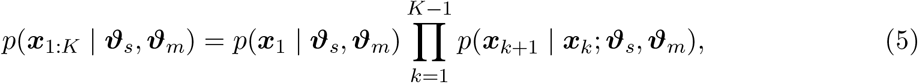

where *p*(***x***_1_ | ***ϑ***_*s*_, ***ϑ***_*m*_) is the prior probability distribution for the allele frequencies of the underlying population at the initial sampling time point, commonly taken to be non-informative (*e.g*., flat over the entire state space) if the prior knowledge is poor, and *p*(***x***_*k*+1_ | ***x***_*k*_; ***ϑ***_*s*_, ***ϑ***_*m*_) is the transition probability density function of the Wright-Fisher diffusion between two consecutive sampling time points for *k* = 1, 2, …, *K* − 1, which satisfies the Kolmogorov backward equation (or its adjoint) resulting from the Wright-Fisher diffusion in Eq. (1). Unless otherwise specified, in this work we take the prior *p*(***x***_1_ | ***ϑ***_*s*_, ***ϑ***_*m*_) to be a uniform distribution over the state space Ω_***X***_, known as the flat Dirichlet distribution, if migration starts before the first sampling time point, *i.e*., *t*_*m*_ ≤ *t*_1_; otherwise, the prior *p*(***x***_1_ | ***ϑ***_*s*_, ***ϑ***_*m*_) is set to be a uniform distribution over the state space Ω_***X***_ restricted to the line satisfying the condition that *x*_3_ = *x*_4_ = 0, *i.e*., there is no continent allele in the island population.

Given the allele frequency trajectories of the underlying population, the observations at each sampling time point are independent. Therefore, we have

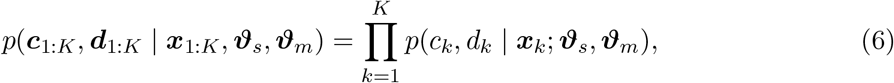

where *p*(*c*_*k*_, *d*_*k*_ | ***x***_*k*_; ***ϑ***_*s*_, ***ϑ***_*m*_) is the probability of the observations at the sampling time point *t*_*k*_ given its corresponding allele frequencies of the underlying population for *k* = 1, 2, …, *K*. If the sample continent allele count *d*_*k*_ is available, we introduce ***z***_*k*_ = (*z*_1,*k*_, *z*_2,*k*_, *z*_3,*k*_, *z*_4,*k*_) to be the counts of the 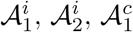 and 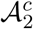 alleles in the sample at the *k*-th sampling time point, and the emission probability *p*(*c*_*k*_, *d*_*k*_ | ***x***_*k*_; ***ϑ***_*s*_, ***ϑ***_*m*_) can be expressed as

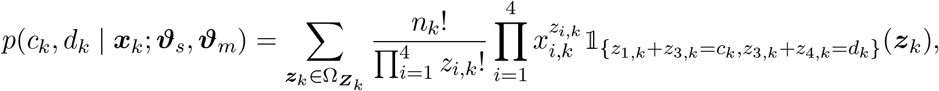

where 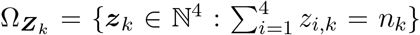 and 𝕝_*A*_ is the indicator function that equals to 1 if condition *A* holds and 0 otherwise. Otherwise, the emission probability *p*(*c*_*k*_, *d*_*k*_ | ***x***_*k*_; ***ϑ***_*s*_, ***ϑ***_*m*_) can be reduced to

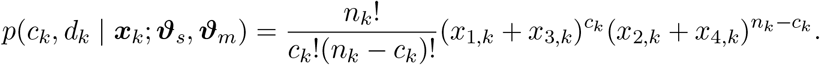

#### 2.2.2. Particle marginal Metropolis-Hastings

The most challenging part in the computation of the posterior *p*(***ϑ***_*s*_, ***ϑ***_*m*_, ***x***_1:*K*_ | ***c***_1:*K*_, ***d***_1:*K*_) is obtaining the transition probability density function *p*(***x***_*k*+1_ | ***x***_*k*_; ***ϑ***_*s*_, ***ϑ***_*m*_) for *k* = 1, 2, …, *K* −1. In principle, the transition probability density function can be achieved by numerically solving the Kolmogorov backward equation (or its adjoint) resulting from the Wright-Fisher diffusion in Eq. (1) typically through a finite difference scheme (Bollback et al., 2008; He et al., 2020c). This requires a fine discretisation of the state space Ω_***X***_ to guarantee numerically stable computation of the solution, but how fine the discretisation needs to be strongly depends on the underlying population genetic quantities that we aim to estimate (He et al., 2020a). We therefore resort to the PMMH algorithm of Andrieu et al. (2010) in this work. The PMMH algorithm only involves simulating the Wright-Fisher SDE in Eq. (1) and permits a joint update of the population genetic parameters and the allele frequency trajectories of the underlying population.

In our PMMH-based procedure, the marginal likelihood

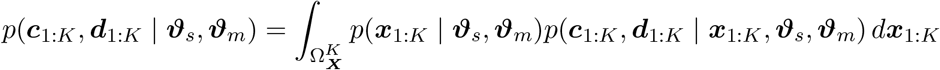

is estimated with the bootstrap particle filter developed by Gordon et al. (1993), where the particles are generated by simulating the Wright-Fisher SDE in Eq. (1) with the Euler-Maruyama scheme. The product of the average weights of the set of particles at the sampling time points ***t***_1:*K*_ yields an unbiased estimate of the marginal likelihood *p*(***c***_1:*K*_, ***d***_1:*K*_ | ***ϑ***_*s*_, ***ϑ***_*m*_), and the underlying population allele frequency trajectories ***x***_1:*K*_ are sampled once from the final set of particles with their corresponding weights. Since the strength of selection and migration can be strongly correlated with their timing, we adopt a blockwise updating scheme to avoid the small acceptance ratio of the PMMH with full dimensional updates. We partition the population genetic parameters into two blocks, the selection-related parameters ***ϑ***_*s*_ and the migration-related parameters ***ϑ***_*m*_, respectively, and iteratively update one block at a time through the PMMH.

More specifically, we first generate a set of the initial candidates of the parameters (***ϑ***_*s*_, ***ϑ***_*m*_) from the prior *p*(***ϑ***_*s*_, ***ϑ***_*m*_). We then run a bootstrap particle filter with the proposed parameters (***ϑ***_*s*_, ***ϑ***_*m*_) to obtain a marginal likelihood estimate 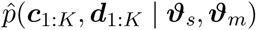 and an initial candidate of the underlying population allele frequency trajectories ***x***_1:*K*_. Repeat the following steps until a sufficient number of the samples of the parameters (***ϑ***_*s*_, ***ϑ***_*m*_) and the underlying population allele frequency trajectories ***x***_1:*K*_ have been obtained:

Step 1: Update the selection-related parameters ***ϑ***_*s*_.

Step 1a: Draw 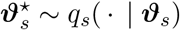.

Step 1b: Run a bootstrap particle filter with 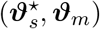 to yield 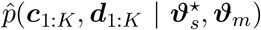 and 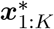.

Step 1c: Accept 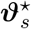 and 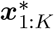 with

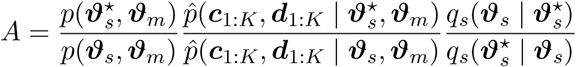

otherwise set 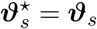 and 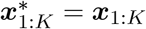.

Step 2: Update the migration-related parameters ***ϑ***_*m*_.

Step 2a: Draw 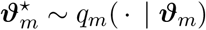.

Step 2b: Run a bootstrap particle filter with 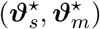 to yield 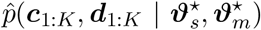 and 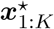.

Step 2c: Accept 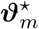 and 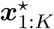 with

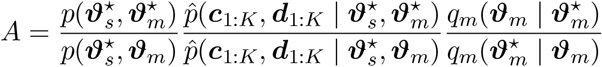

otherwise set 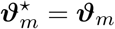 and 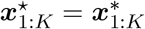.

In this work we use random walk proposals for both selection- and migration-related parameters in our blockwise PMMH algorithm unless otherwise specified.

Once enough samples of the parameters (***ϑ***_*s*_, ***ϑ***_*m*_) and the underlying population allele frequency trajectories ***x***_1:*K*_ have been yielded, we can compute the posterior *p*(***ϑ***_*s*_, ***ϑ***_*m*_ | ***c***_1:*K*_, ***d***_1:*K*_) from the samples of the parameters (***ϑ***_*s*_, ***ϑ***_*m*_) using nonparametric density estimation techniques (see Izenman, 1991, for a review) and achieve the maximum a posteriori probability (MAP) estimates for the population genetic parameters. Our estimates for the underlying population allele frequency trajectories are the posterior mean of the stored samples of the underlying population allele frequency trajectories. Our approach can be readily extended to the analysis of multiple (independent) loci. Given that the migration-related parameters are shared by all loci, in each iteration we only need to replicate Step 1 once to update selection-related parameters specified for each locus and then update migration-related parameters with shared by all loci in Step 2, where the likelihood is replaced by the product of the likelihoods for each locus.

## 3. Results

In this section, we first evaluate the performance of our approach using simulated datasets with various population genetic parameter values and then apply it to re-analyse the time series allele frequency data from ancient chicken in Loog et al. (2017) genotyped at the locus encoding for the thyroid-stimulating hormone receptor (*TSHR*), which is hypothesised to have undergone strong and recent selection in domestic chicken.

### 3.1. Robustness and performance

To assess our method, we ran forward-in-time simulations of the multi-allele Wright-Fisher model with selection and migration described in File S1 and examined the bias and the root mean square error (RMSE) of our estimates obtained from these replicate simulations. Here we varied the selection coefficient *s* ∈ {0.003, 0.006, 0.009} and the dominance parameter *h* ∈ {0, 0.5, 1} and fixed the migration rate *m* = 0.005. We adopted the selection time *k*_*s*_ = 180 and varied the migration time *k*_*m*_ ∈ {90, 360}, which were measured in generations. In addition, we varied the population size *N* ∈ {5000, 50000, 500000}. In principle, the conclusions we draw here hold for other values of the population genetic parameters in similar ranges.

More specifically, we ran two groups of simulation. For the first group, we fix the dominance parameter *h* = 0.5 and vary all other parameters in the sets specified above, yielding a total of 18 parameter combinations. For the second group, we fix the population size *N* = 50000 and vary all other parameters, yielding another 18 parameter combinations. Due to overlap between these two groups, we have a total of 30 parameter combinations, for each of which we performed 300 replicated runs. For each run, we took the starting allele frequencies of the underlying island population at generation 0 (*i.e*., the first sampling time point) to be ***x***_1_ = (0.4, 0.6, 0, 0) and the mutant allele frequency of the underlying continent population to be *x*_*c*_ = 0.9. These values are similar to those of ancient chicken samples reported in Loog et al. (2017). We simulated a total of 500 generations under the multi-allele Wright-Fisher model with selection and migration and generated a multinomial sample of 100 chromosomes from the underlying island population every 50 generations from generation 0, 11 sampling time points in total. At each sampling time point, we generated the mutant allele count by summing the first and third components of the simulated sample allele counts, and the continent allele count by summing the third and fourth components since in real data only mutant allele counts and continent allele counts are available. Additionally, in real data the continent allele count of the sample may not be available at each sampling time point (*e.g*., Loog et al., 2017). To explore the impact of missing continent allele counts, we assumed that the continent allele counts of the sample were unavailable at the first three and seven sampling time points, respectively, for each run in our simulation studies, as shown in simulated datasets A and B, respectively, in Figure 1.

**Figure 1:**
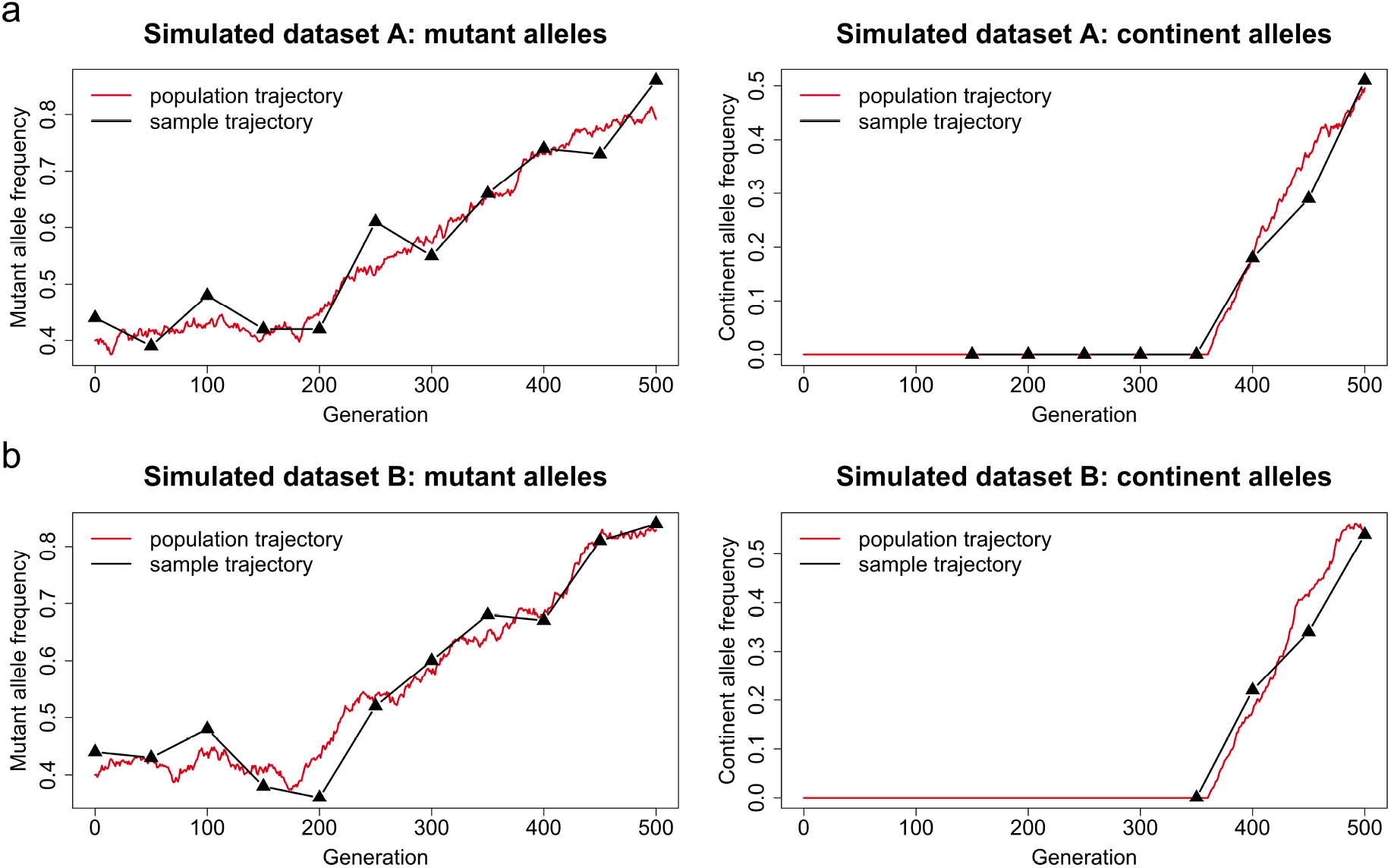
Representative examples of the datasets simulated using the Wright-Fisher model with selection and migration. We take the selection coefficient and time to be *s* = 0.006 and *k*_*s*_ = 180 and the migration rate and time to be *m* = 0.005 and *k*_*m*_ = 360, respectively. We set the dominance parameter *h* = 0.5 and the population size *N* = 5000. We adopt the starting allele frequencies of the underlying island population ***x***_1_ = (0.4, 0.6, 0, 0) and the mutant allele frequency of the underlying continent population *x*_*c*_ = 0.9. We sample 100 chromosomes at every 50 generations from generation 0 to 500. (a) simulated dataset A: continent allele counts are not available at the first three sampling time points. (b) simulated dataset B: continent allele counts of the sample are not available at the first seven sampling time points.

In our procedure, we chose a uniform prior over the interval [−1, 1] for the selection coefficient *s* and a uniform prior over the interval [0, 1] for the migration rate *m*. We let the starting times of selection and migration *k*_*s*_ and *k*_*m*_ be uniformly distributed over the set of all possible time points, *i.e*., *k*_*s*_, *k*_*m*_ ∈ {*k* ∈ ℤ: *k* ≤ 500}. We generated 10000 iterations of the blockwise PMMH with 1000 particles, and in the Euler-Maruyama scheme each generation was divided into five subintervals. We discarded the first half of the iterations as the burn-in period and then thinned the remaining PMMH output by selecting every fifth value. See Figures 2 and 3 for our posteriors for the timing and strength of selection and migration based on the simulated datasets shown in Figure 1, including our estimates for the mutant and continent allele frequency trajectories of the underlying island population. Evidently, our approach is capable of identifying the selection and migration signatures and accurately estimate their timing and strength in these two examples. The underlying frequency trajectories of the mutant and continent alleles in island population are both well matched with our estimates, *i.e*., the allele frequency trajectories of the underlying island population fluctuate slightly around our estimates and are completely covered by our 95% highest posterior density (HPD) intervals.

**Figure 2:**
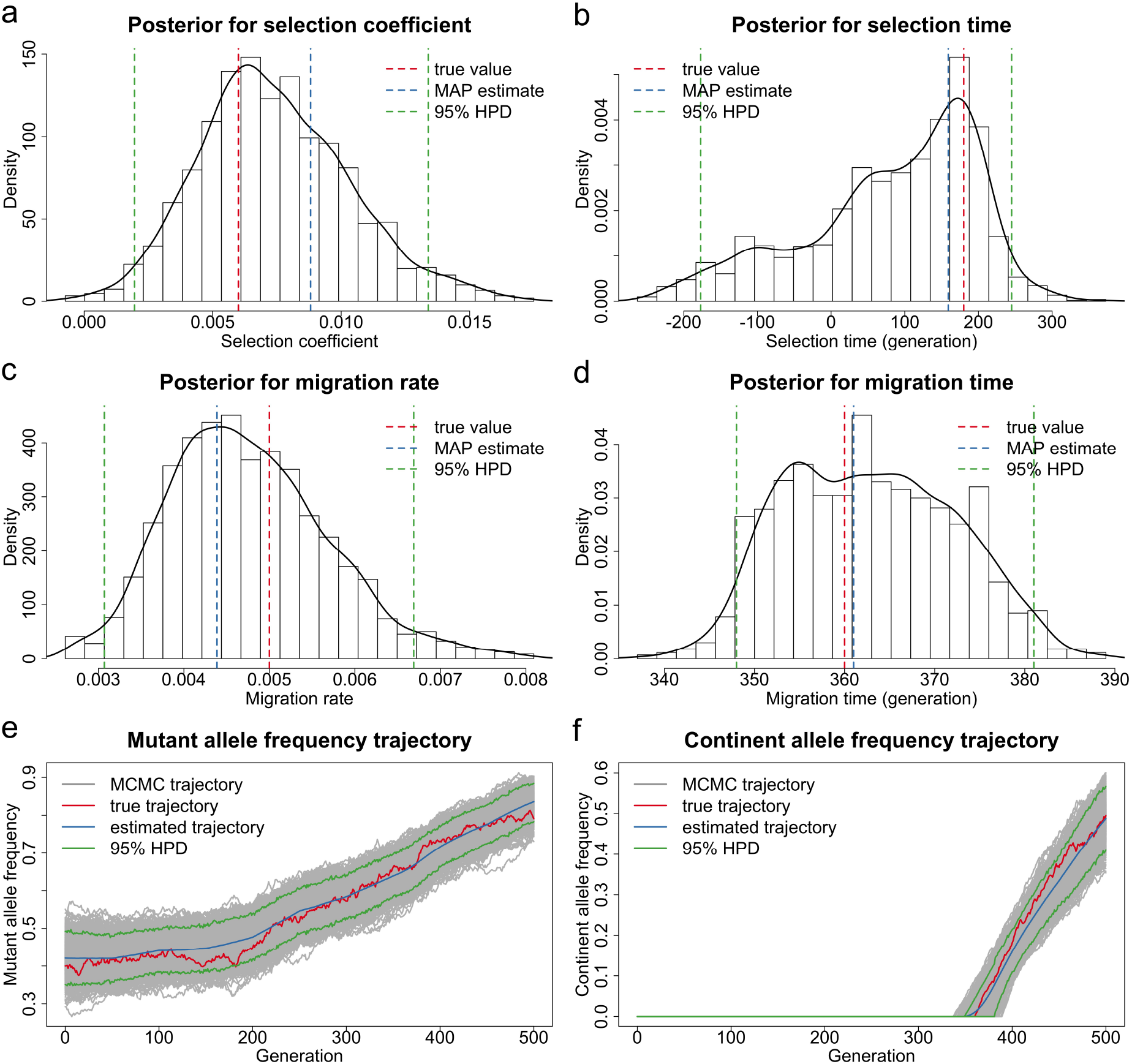
Bayesian estimates for the dataset shown in Figure 1a simulated for the case of the continent allele counts unavailable at the first three sampling time points. Posteriors for (a) the selection coefficient (b) the selection time (c) the migration rate and (d) the migration time. The MAP estimate is for the joint posterior, and may not correspond to the mode of the marginals. Estimated underlying trajectories of (e) the mutant allele frequency and (f) the continent allele frequency of the island population.

**Figure 3:**
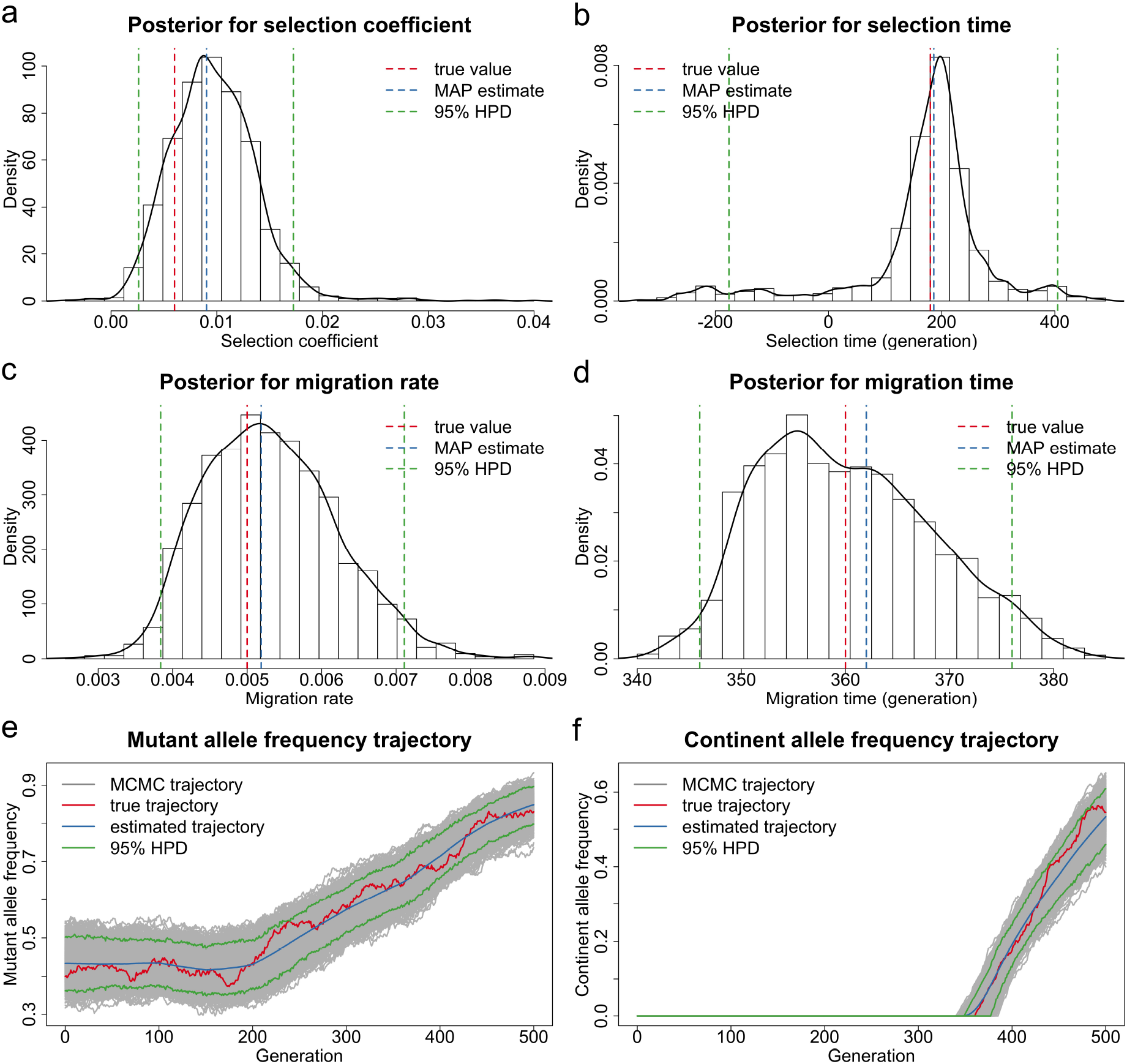
Bayesian estimates for the dataset shown in Figure 1b simulated for the case of continent allele counts unavailable at the first seven sampling time points. Posteriors for (a) the selection coefficient (b) the selection time (c) the migration rate and (d) the migration time. The MAP estimate is for the joint posterior, and may not correspond to the mode of the marginals. Estimated underlying trajectories of (e) the mutant allele frequency and (f) the continent allele frequency of the island population.

In Figure 4, we present the boxplots of our estimates for additive selection (*h* = 0.5) where continent allele counts are not available at the first three sampling time points. These boxplots show the relative bias of (a) the selection coefficient estimates, (b) the selection time estimates, (c) the migration rate estimates and (d) the migration time estimates across 18 different combinations of the selection coefficient, the migration time and the population size. The tips of the whiskers represent the 2.5%-quantile and the 97.5%-quantile, and the boxes denote the first and third quartiles with the median in the middle. We summarise the bias and the RMSE of the estimates in Tables S1 and S2.

**Figure 4:**
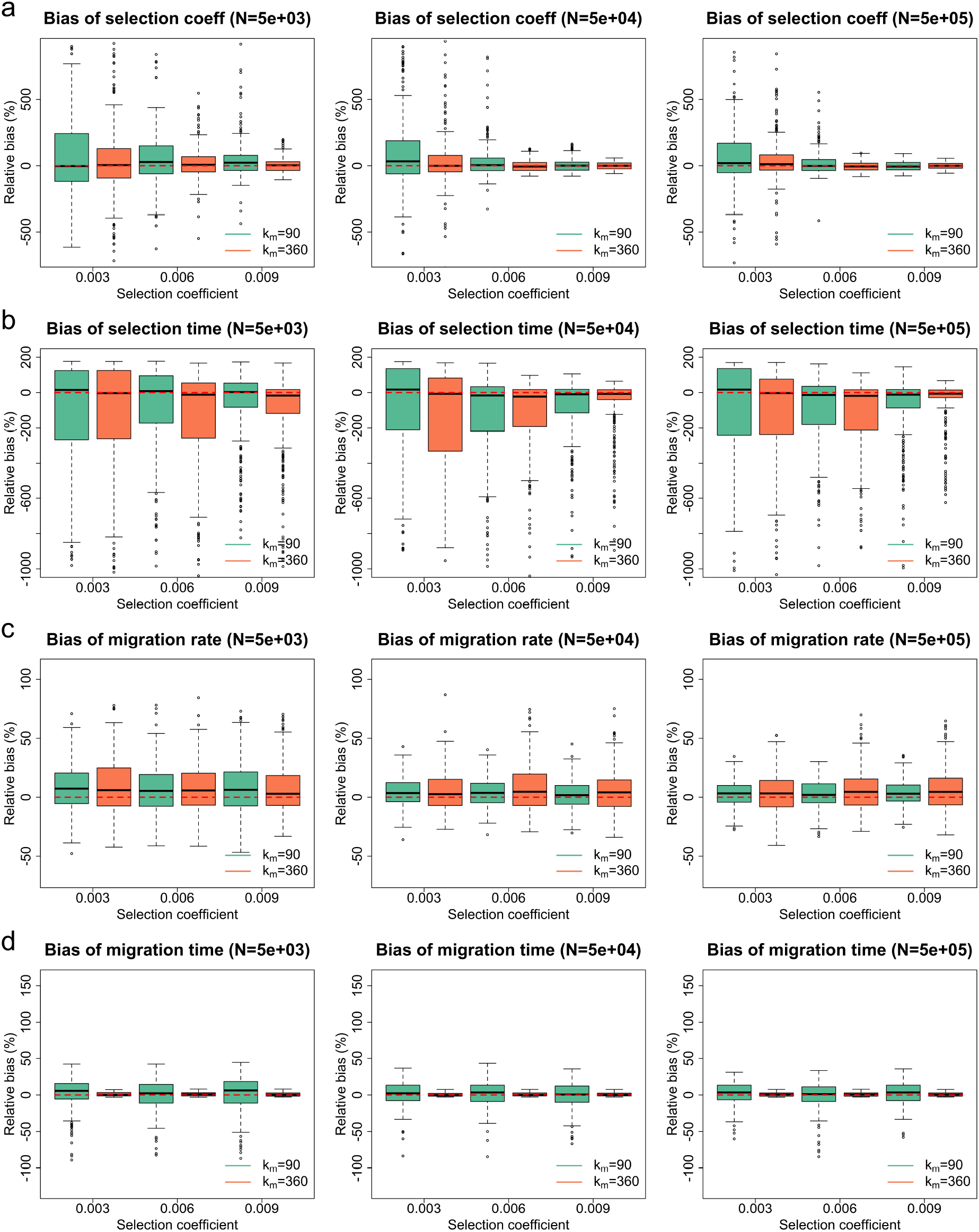
Empirical distributions of the estimates for 300 datasets simulated for additive selection (*h* = 0.5) where continent allele counts are not available at the first three sampling time points. Green boxplots represent the estimates produced for the case of selection starting after migration, and orange boxplots represent the estimates produced for the case of selection starting before migration. Boxplots of the relative bias of (a) the selection coefficient estimates (b) the selection time estimates (c) the migration rate estimates and (d) the migration time estimates. To aid visual comparison, we have picked the *y* axes here so that boxes are of a relatively large size. This causes some outliers to lie outside the plots. Boxplots containing all outliers can be found in Figure S1.

As shown in Figure 4, our estimates for the selection coefficient and time are approximately median-unbiased across 18 different parameter combinations, but the migration rate and time are both slightly overestimated (*i.e*., a small positive bias is found in our estimates). An increase in the population size results in the better overall performance of our estimation (*i.e*., smaller bias with smaller variance). In particular, the average proportion of the replicates for which the signature of selection can be identified (*i.e*., the 95% HPD interval does not contain the value of 0) increases from 17.17% to 59.33% and then to 80.67% as the population size increases. Such an improvement in the performance of our estimation is to be expected since large population sizes reduce the magnitude of the stochastic effect on the changes in allele frequencies due to genetic drift, which dilutes evidence of selection and migration.

Compared to the case of selection starting after migration (*i.e*., *k*_*m*_ = 90), our estimates for the case of selection starting before migration (*i.e*., *k*_*m*_ = 360) reveal smaller bias and variances in both selection coefficient and time. The average proportion of replicates where the signature of selection can be identified when the migration time *k*_*m*_ = 360 is 15.96% higher than when *k*_*m*_ = 90. One possible explanation is as follows: if selection begins before migration, there is a period of time that the allele frequency trajectories of the underlying population are only under the influence of selection. In contrast, our method performs better for the migration rate when selection starts after migration (*i.e*., *k*_*m*_ = 90), but the performance for the migration time deteriorates somewhat unexpectedly when the migration time *k*_*m*_ = 360. This might be due to our parameter setting where the starting time of migration is within the period of availability of continent allele counts for *k*_*m*_ = 360, but not for *k*_*m*_ = 90.

In addition, we see from Figure 4 that the bias and variance of our estimates for the selection coefficient and time are largely reduced as the selection coefficient increases, especially in terms of outliers. The average proportion of the replicates where selection signatures can be identified increases from 27.56% to 63.11% and then to 66.50% as the selection coefficient increases, with 97.17% for the case of large population size (*N* = 500000) and selection coefficient (*s* = 0.009). For weak selection, the underlying trajectory of allele frequencies is extremely stochastic so that it is difficult to disentangle the effects of genetic drift and natural selection (Schraiber et al., 2013). An increase in the strength of selection leads to more pronounced changes through time in allele frequencies, making the signature of selection more identifiable. In contrast, an increase in the selection coefficient has little effect on our estimates of the migration rate and time.

In Figure 5, we present the boxplots of our estimates for additive selection (*h* = 0.5) where continent allele counts are unavailable at the first seven sampling time points, with their bias and RMSE summarised in Tables S3 and S4. They reveal similar behaviour in estimation bias and variance, although our estimates for the migration-related parameters show significantly larger bias and variances, probably resulting from the increased length of time when continent allele counts are not available. This, however, has little effect on our estimation of the selection-related parameters, with similar average proportions of the replicates where the signature of selection can be identified (52.39% *vs*. 52.17%).

**Figure 5:**
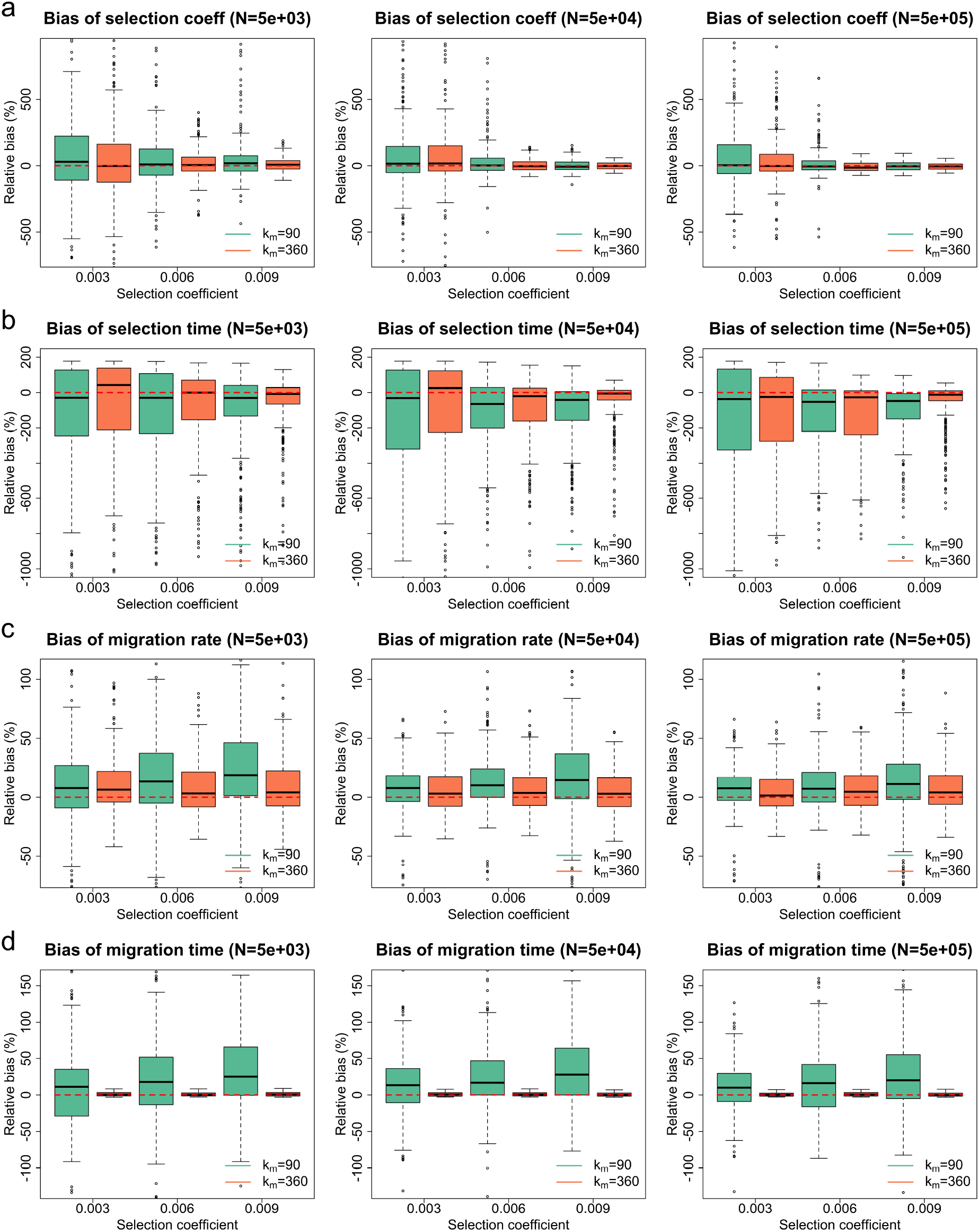
Empirical distributions of the estimates for 300 datasets simulated for additive selection (*h* = 0.5) where continent allele counts are not available at the first seven sampling time points. Green boxplots represent the estimates produced for the case of selection starting after migration, and orange boxplots represent the estimates produced for the case of selection starting before migration. Boxplots of the relative bias of (a) the selection coefficient estimates (b) the selection time estimates (c) the migration rate estimates and (d) the migration time estimates. To aid visual comparison, we have picked the *y* axes here so that boxes are of a relatively large size. This causes some outliers to lie outside the plots. Boxplots containing all outliers can be found in Figure S2.

The resulting estimates for dominant selection (*h* = 0) and recessive selection (*h* = 1) can be found in Figures S3 and S4, respectively, for the case of the population size *N* = 50000. They are very similar to the boxplot results in the empirical studies for additive selection illustrated in Figures 4 and 5. Their bias and RMSE are summarised in Tables S5-S8. It should be noted that overall, recessive selection yields the best performance, additive selection next, while dominant selection yields the worst performance in our simulation studies for the inference of selection. This is mainly due to our parameter setting, *i.e*., the effect of selection, when the mutant allele has been *established* in the population (*e.g*., our starting mutant allele frequency is 0.4), is the strongest for recessive selection and weakest for dominant selection (see Figure S5).

In conclusion, our approach can produce reasonably accurate joint estimates of the timing and strength of selection and migration from time series data of allele frequencies across different parameter combinations. Our estimates for the selection coefficient and time are approximately median-unbiased, with smaller variances as the population size or the selection coefficient (or both) increases. Our estimates for the migration rate and time both show little positive bias. Their performance improves with an increase in population size or the number of the sampling time points when continent allele counts are available (or both).

### 3.2. Application to ancient chicken samples

We re-analysed aDNA data of 452 European chicken genotyped at the *TSHR* locus (position 43250347 on chromosome 5) from previous studies of Flink et al. (2014) and Loog et al. (2017). The time from which the data come ranges from approximately 2200 years ago to the present. The data shown in Table 2 come from grouping the raw chicken samples by their sampling time points. The raw sample information and genotyping results can be found in Loog et al. (2017). The derived *TSHR* allele has been associated with reduced aggression to conspecifics and faster onset of egg laying (Belyaev, 1979; Rubin et al., 2010; Karlsson et al., 2015,2016), which was hypothesised to have undergone strong and recent selection in domestic chicken (Rubin et al., 2010; Karlsson et al., 2015) from the period of time when changes in Medieval dietary preferences and husbandry practices across northwestern Europe occurred (Loog et al., 2017).

**Table 2:** Time serial European chicken samples of segregating alleles at the *TSHR* locus. The unit of the sampling time is the AD year so that positive values denote the AD year *e.g*., AD 82, and negative values denote the BC year, *e.g*., 25 BC.

| Sample time | Sample size | Mutant allele |
| --- | --- | --- |
| -128 | 12 | 8 |
| -25 | 8 | 5 |
| 82 | 8 | 3 |
| 200 | 32 | 14 |
| 256 | 14 | 3 |
| 1067 | 6 | 0 |
| 1309 | 20 | 18 |
| 1650 | 2 | 1 |
| 1850 | 2 | 2 |
| 1975 | 14 | 14 |
| 1995 | 334 | 328 |

To avoid overestimating the effect of selection on allele frequency changes, we model recent migration in domestic chicken from Asia to Europe in this work. More specifically, the European chicken population was represented as the island population while the Asian chicken population was represented as the continent population with a derived *TSHR* allele frequency of *x*_*c*_ = 0.99 fixed from the time of the onset of migration, which is a conservative estimate chosen in Loog et al. (2017). Migration from Asia in domestic chicken, beginning around 250 years ago and continuing until the present, was historically well documented (Dana et al., 2011; Flink et al., 2014; Lyimo et al., 2015). Unlike Loog et al. (2017), we estimated the migration rate along with the selection coefficient and time by incorporating the estimate reported in Loog et al. (2017) that about 15% of the modern European chicken have Asia origin. This allows us to obtain the sample frequency of the allele in European chicken at the most recent sampling time point that resulted from immigration from Asia. We took the average length of a generation of chicken to be one year, and the time measured in generations was offset so that the most recent sampling time point was generation 0.

In our analysis, we adopted the dominance parameter *h* = 1 since the derived *TSHR* allele is recessive, and picked the population size *N* = 180000 (95% HPD 26000-460000) estimated by Loog et al. (2017). We chose a uniform prior over the interval [−1, 1] for the selection coefficient *s* and a uniform prior over the set {−9000, −8999, …, 0} for the selection time *k*_*s*_, which covers chicken domestication dated to about 8000 (95% CI 7014–8768) years ago (Lawal et al., 2020). We picked a uniform prior over the interval [0, 1] for the migration rate *m* and set the migration time *k*_*m*_ = −250. All settings in the Euler-Maruyama scheme and the blockwise PMMH algorithm are the same as we applied in Section 3.1. The posteriors for the selection coefficient, the selection time and the migration rate are illustrated in Figure 6, as well as the estimates for the underlying frequency trajectories of the mutant and Asian alleles in the European chicken population. The MAP estimates, as well as 95% HPD intervals, are summarised in Table 3.

**Figure 6:**
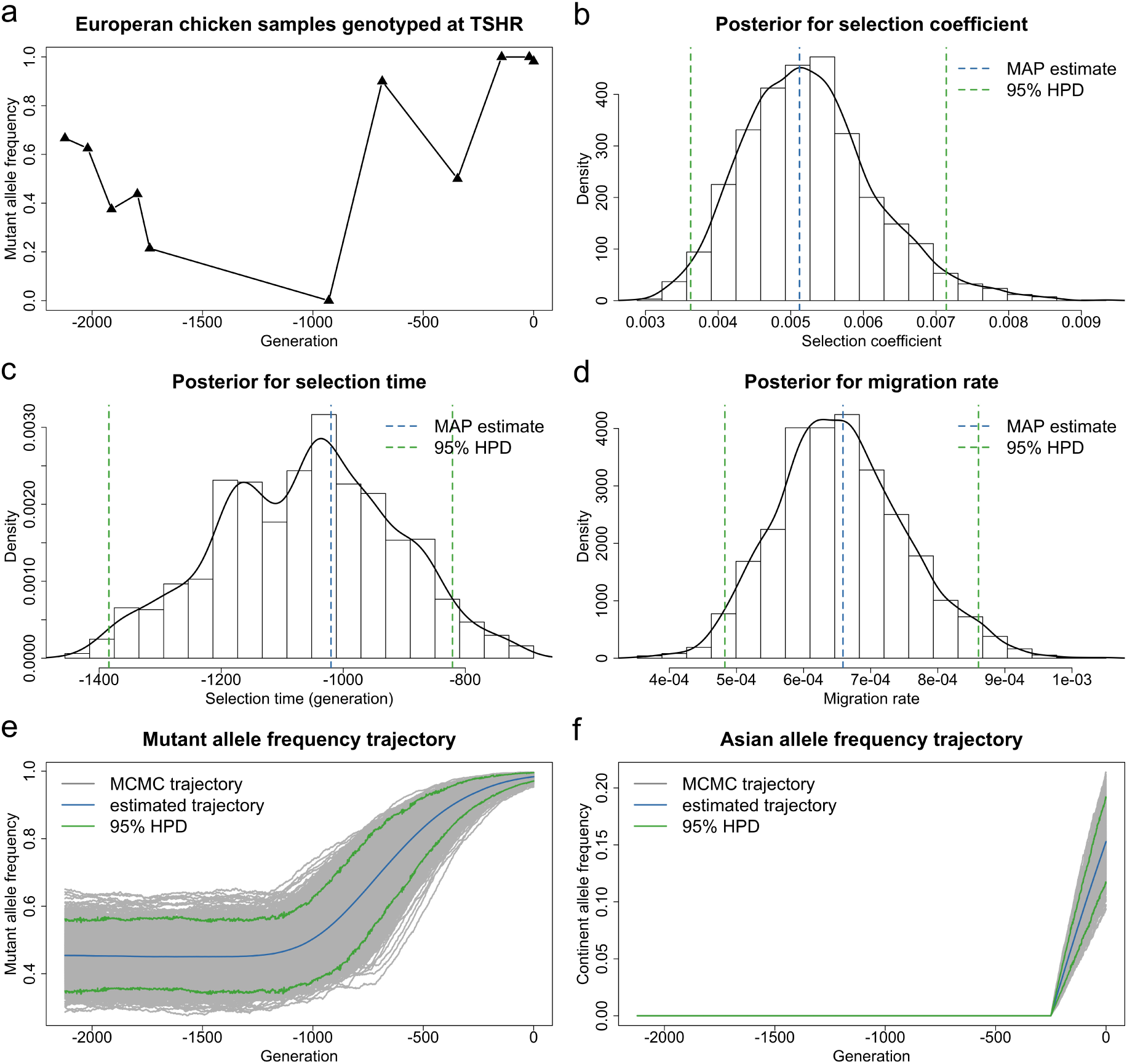
Bayesian estimates for aDNA data of European chicken genotyped at the *TSHR* locus from Loog et al. (2017) for the case of the population size *N* = 180000. (a) Temporal changes in the mutant allele frequencies of the sample, where the sampling time points have been offset so that the most recent sampling time point (AD 1995) is generation 0. Posteriors for (b) the selection coefficient (c) the selection time and (d) the migration rate. Estimated underlying trajectories of (e) the mutant allele frequency and (f) the Asian allele frequency in the European chicken population. The MAP estimate is for the joint posterior, and may not correspond to the mode of the marginals.

**Table 3:**
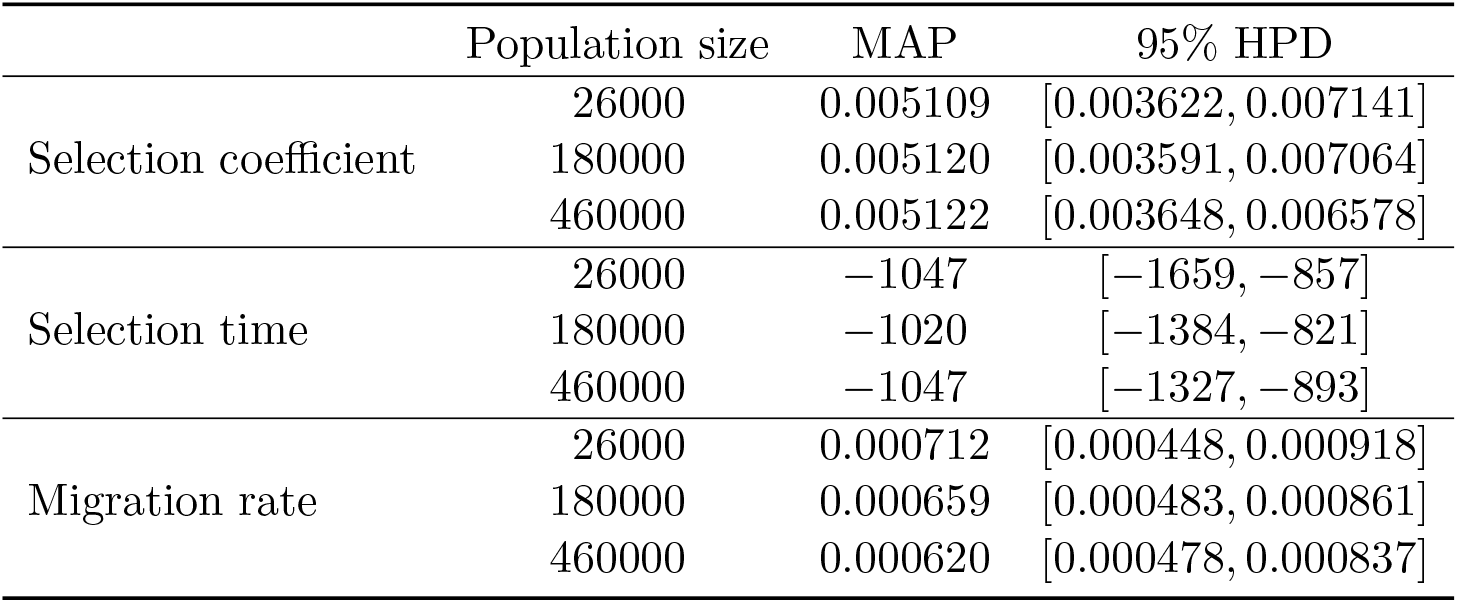
MAP estimates of the selection coefficient, the selection time and the migration rate, as well as their 95% HPD intervals, for *TSHR* achieved with the population size *N* = 26000, *N* = 180000 and *N* = 460000.

|  | Population size | MAP | 95% HPD |
| --- | --- | --- | --- |
| Selection coefficient | 26000 | 0.005109 | [0.003622, 0.007141] |
|  | 180000 | 0.005120 | [0.003591, 0.007064] |
|  | 460000 | 0.005122 | [0.003648, 0.006578] |
| Selection time | 26000 | -1047 | [-1659, -857] |
|  | 180000 | -1020 | [-1384, -821] |
|  | 460000 | -1047 | [-1327, -893] |
| Migration rate | 26000 | 0.000712 | [0.000448, 0.000918] |
|  | 180000 | 0.000659 | [0.000483, 0.000861] |
|  | 460000 | 0.000620 | [0.000478, 0.000837] |

From Table 3, we observe that our estimate of the selection coefficient for the mutant allele is 0.005120 with 95% HPD interval [0.003591, 0.007064], strong evidence to support the derived *TSHR* allele being selectively advantageous in the European chicken population. Such positive selection results in an increase over time in the mutant allele frequency, starting from AD 975 with 95% HPD interval [611, 1174] (see Figure 6e). The starting frequency of the derived *TSHR* allele in 128 BC is 0.454200 with 95% HPD interval [0.349024, 0.562094], which is similar to that estimated in a red junglefowl captive zoo population in Rubin et al. (2010). Our estimate of the migration rate for the Asian allele is 0.000659 with 95% HPD interval [0.000483, 0.000861]. This migration, starting about 250 years ago, leads to 15.2848% of the European chicken with Asian ancestry in AD 1995, with 95% HPD interval [0.116412, 0.191382] (see Figure 6f). Our findings are consistent with those reported in Loog et al. (2017). This is further confirmed by the results obtained with different values of the population size (*i.e*., *N* = 26000 and *N* = 460000, the lower and upper bounds of 95% HPD interval for the population size given in Loog et al. (2017), respectively). These results are shown in Figures S6 and S7 and summarised in Table 3.

To evaluate the performance of our approach when samples are sparsely distributed in time with small uneven sizes like the European chicken samples at the *TSHR* locus we have studied above, we generated 300 simulated datasets that mimic the *TSHR* data, *i.e*., we used the sample times and sizes as given in Table 2, the timing and strength of selection and migration as given by MAP estimates found in Table 3, and population size *N* = 180000. From Figure 7, we find that our simulation studies based on the *TSHR* data yield median-unbiased estimates for the selection coefficient, the selection time and the migration rate, similar to our performance in the simulation studies shown in Section 3.1. Moreover, the signature of selection can be identified in all 300 replicates. This illustrates that our method can achieve good performance even though samples are sparsely distributed in time with small uneven sizes, which is highly desirable for aDNA data.

**Figure 7:**
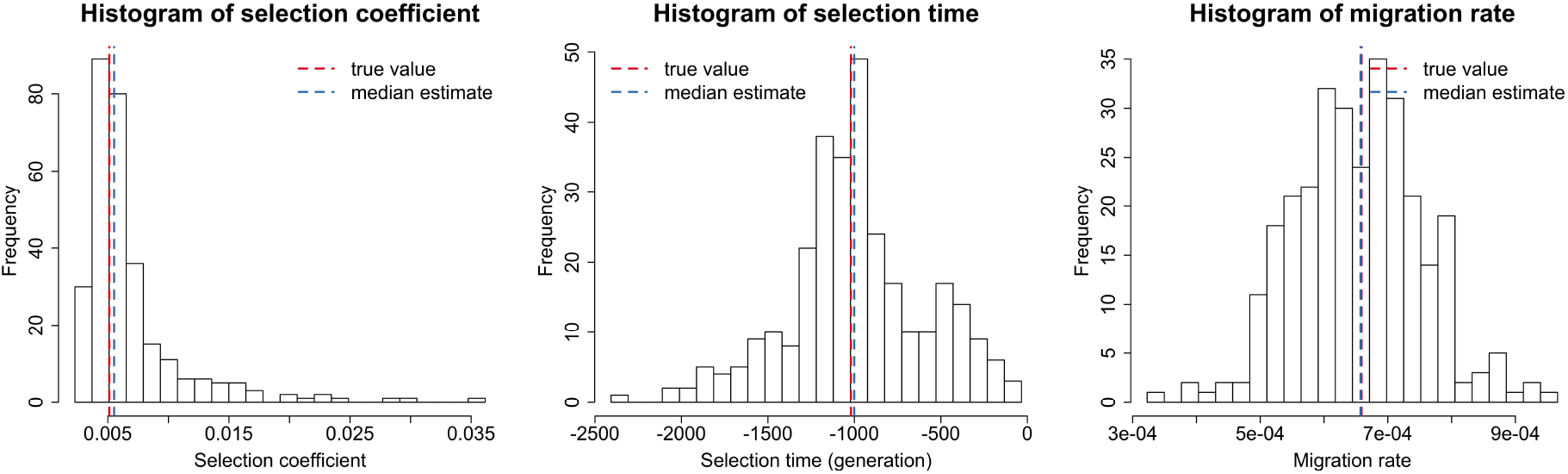
Empirical distributions of the estimates for 300 datasets simulated for *TSHR* based on the aDNA data shown in Table 2. We simulate the underlying population dynamics with the timing and strength of selection and migration estimated with the population size *N* = 180000 shown in Table 3. To aid visual comparison, we have picked the *x* axis in the left panel not to cover all 300 estimates. The histogram containing all 300 estimates can be found in Figure S8.

In summary, our approach works well on the ancient chicken samples, even though they are sparsely distributed in time with small uneven sizes. Our estimates demonstrate strong evidence for the derived *TSHR* allele being positively selected between the 7th and 12th centuries, which coincides with the time period of changes in dietary preferences and husbandry practices across northwestern Europe. This again shows possible links established by Loog et al. (2017) between the selective advantage of the derived *TSHR* allele and a historically attested cultural shift in food preference in Medieval Europe.

## 4. Discussion

In this work, we introduced a novel MCMC-based procedure for the joint inference of the timing and strength of selection and migration from aDNA data. To our knowledge, Mathieson & McVean (2013) and Loog et al. (2017) described the only existing methods that can jointly infer selection and migration from time series data of allele frequencies. However, the approach of Mathieson & McVean (2013) cannot estimate the time of the onset of selection and migration. Loog et al. (2017) only showed the applicability of their approach in the scenario where timing and strength of migration were both pre-specified. In addition, their method is restricted by the assumption of infinite population size, which limits the application of their approach to aDNA. Our method was built upon an HMM framework incorporating a multi-allele Wright-Fisher diffusion with selection and migration. Our estimates for the timing and strength of selection and migration were obtained by the PMMH algorithm with blockwise sampling, which enables the co-estimation of the underlying trajectories of allele frequencies through time as well. This is a highly desirable feature for aDNA because it allows us to infer the drivers of selection and migration by correlating genetic variation patterns with potential evolutionary events such as changes in the ecological context in which an organism has evolved.

We showed through extensive simulation studies that our method could deliver reasonably accurate estimates for the timing and strength of selection and migration, including the estimates for the underlying trajectories of allele frequencies through time. The estimates for the selection coefficient and time were largely unbiased, while the estimates for the migration rate and time showed a slight positive bias. We applied our approach to re-analyse ancient European chicken samples genotyped at the *TSHR* locus from earlier studies of Flink et al. (2014) and Loog et al. (2017). We observed that the derived *TSHR* allele became selectively advantageous from AD 975 (95% HPD 611-1174), which was similar to that reported in Loog et al. (2017). Our results further confirmed the findings of Loog et al. (2017) that positive selection acting on the derived *TSHR* allele in European chicken could be driven by chicken intensification and egg production in Medieval Europe as a result of Christian fasting practices (*i.e*., the consumption of birds, eggs and fish became allowed (Venarde, 2011)). Except for religiously inspired dietary preferences, this could also be a result of changes in Medieval husbandry practices along with population growth and urbanisation in the High Middle Ages (around AD 1000-1250). See Loog et al. (2017) and references cited therein for more details.

Unlike Loog et al. (2017), our approach models genetic drift. From Table 3, we observe that our estimates from aDNA data for *TSHR* are close to each other regardless of what population size we choose from the 95% HPD interval for the European chicken population size reported in Loog et al. (2017). This indicates that ignoring genetic drift might have little effect on the inference of selection from aDNA data like those in Loog et al. (2017). To further investigate the effect of genetic drift, we simulated 300 datasets based on the aDNA data for *TSHR*, where the timing and strength of selection and migration were taken to be our estimates given in Table 3 but the true population size was taken to be *N* = 4500. We ran our method with a misspecified population size *N* = 180000 for these 300 replicates and find that this larger population size leads to significant overestimation of the selection coefficient and time with much larger variance (see Figure 8), which implies the necessity of modelling genetic drift in the inference of selection from aDNA data.

**Figure 8:**
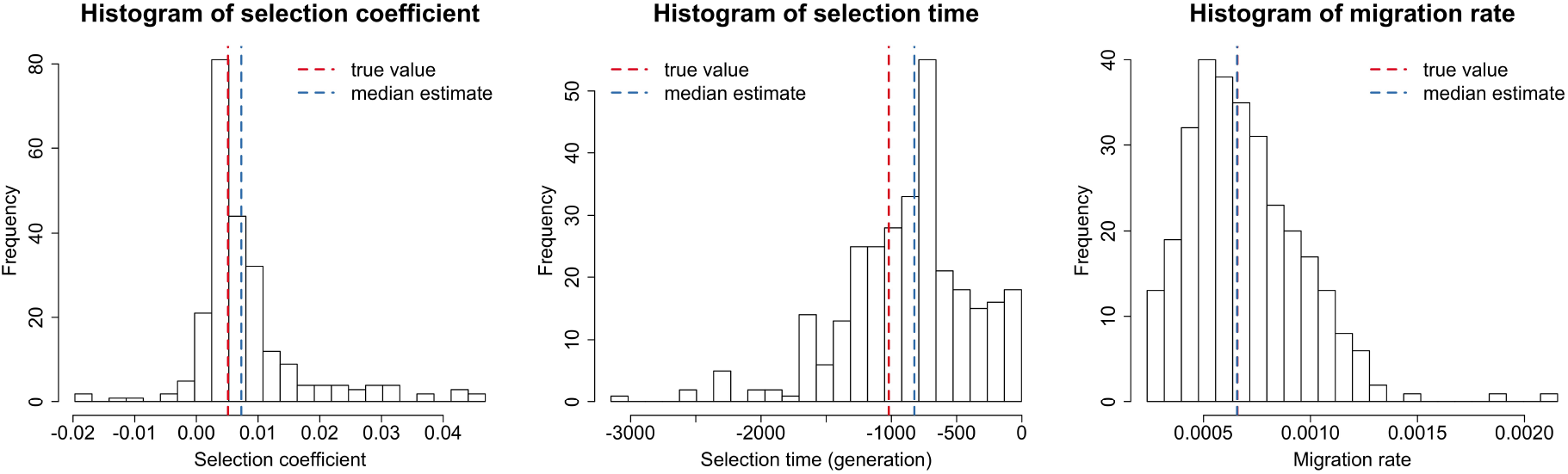
Empirical distributions of the estimates for 300 datasets simulated for *TSHR* based on the aDNA data presented in Table 2. We take the timing and strength of selection and migration to be those estimated with the population size *N* = 180000 given in Table 3, but the true population size in the simulation is taken to be *N* = 4500. To aid visual comparison, we have picked the *x* axis in the left panel not to cover all 300 estimates. The histogram containing all 300 estimates can be found in Figure S9.

We explored how misspecification of genetic dominance or gene migration affects our inference of selection and migration in a similar way. We first simulated 300 datasets based on the aDNA data for *TSHR* with the dominance parameter *h* = 0 and *h* = 0.5, respectively, but we ran our inference procedure with a misspecified dominance parameter *h* = 1. As shown in Figure 9, we find that a misspecified dominance parameter introduces certain bias in the inference results for both selection and migration. We then simulated 300 datasets based on the aDNA data for *TSHR* with the migration rate *m* = 0.00001 and *m* = 0.01, respectively, but we ran our procedure with a misspecified migration rate *m* = 0.000659 (*i.e*., the migration rate estimated with the population size *N* = 180000). We observe from Figure 10 that a misspecified migration rate does not dramatically alter the posterior median of the selection coefficient and time but significantly increase the variance of their estimates. Finally, we simulated 300 datasets based on the aDNA data for *TSHR* with the migration time *k*_*m*_ = −400 and *k*_*m*_ = −100, respectively, but we ran our procedure with a misspecified migration time *k*_*m*_ = −250. From Figure 11, we see that a misspecified migration time has little effect on the inference of selection but dramatically alter the estimate of the migration rate. All these results show the necessity of the joint inference of selection and migration from aDNA data.

**Figure 9:**
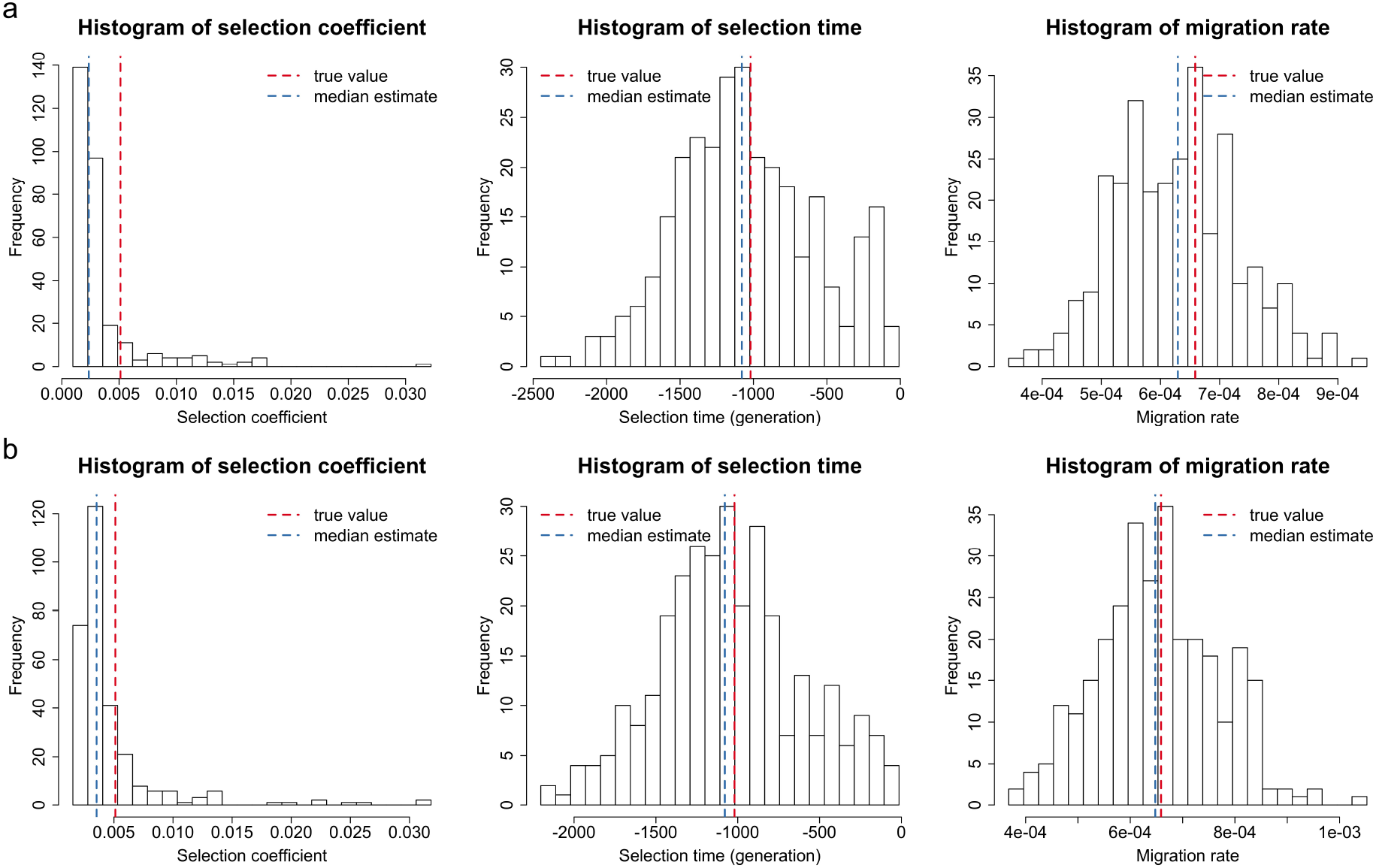
Empirical distributions of the estimates for 300 datasets simulated for *TSHR* based on the aDNA data presented in Table 2. We take the timing and strength of selection and migration to be those estimated with the population size *N* = 180000 given in Table 3, but the true dominance parameter in the simulation is taken to be (a) *h* = 0 and (b) *h* = 0.5, respectively. To aid visual comparison, we have picked the *x* axis in the left panel not to cover all 300 estimates. The histogram containing all 300 estimates can be found in Figure S10.

**Figure 10:**
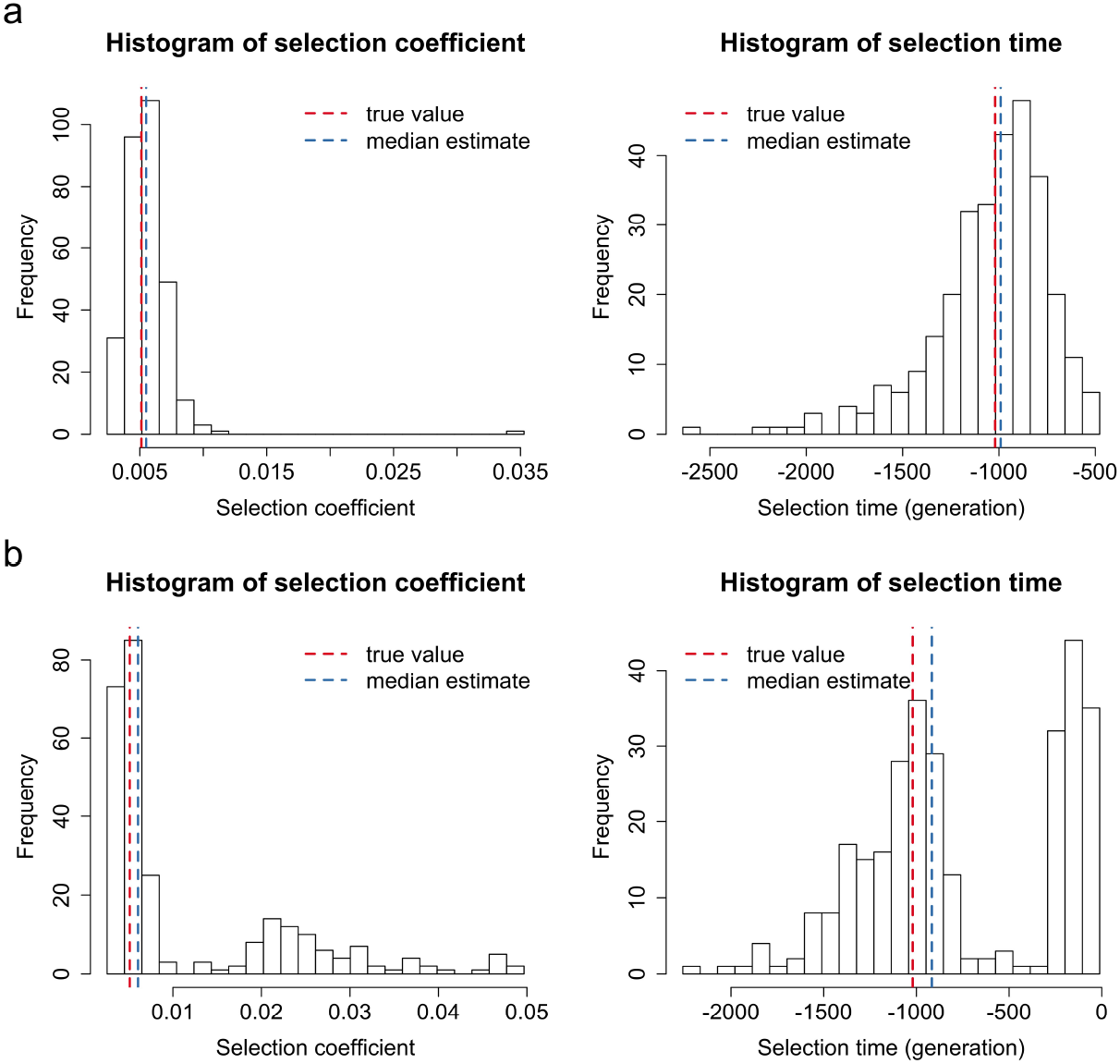
Empirical distributions of the estimates for 300 datasets simulated for *TSHR* based on the aDNA data presented in Table 2. We take the timing and strength of selection and migration to be those estimated with the population size *N* = 180000 given in Table 3, but the true migration rate in the simulation is taken to be (a) *m* = 0.0001 and (b) *m* = 0.001, respectively. To aid visual comparison, we have picked the *x* axis in the left panel not to cover all 300 estimates. The histogram containing all 300 estimates can be found in Figure S11.

**Figure 11:**
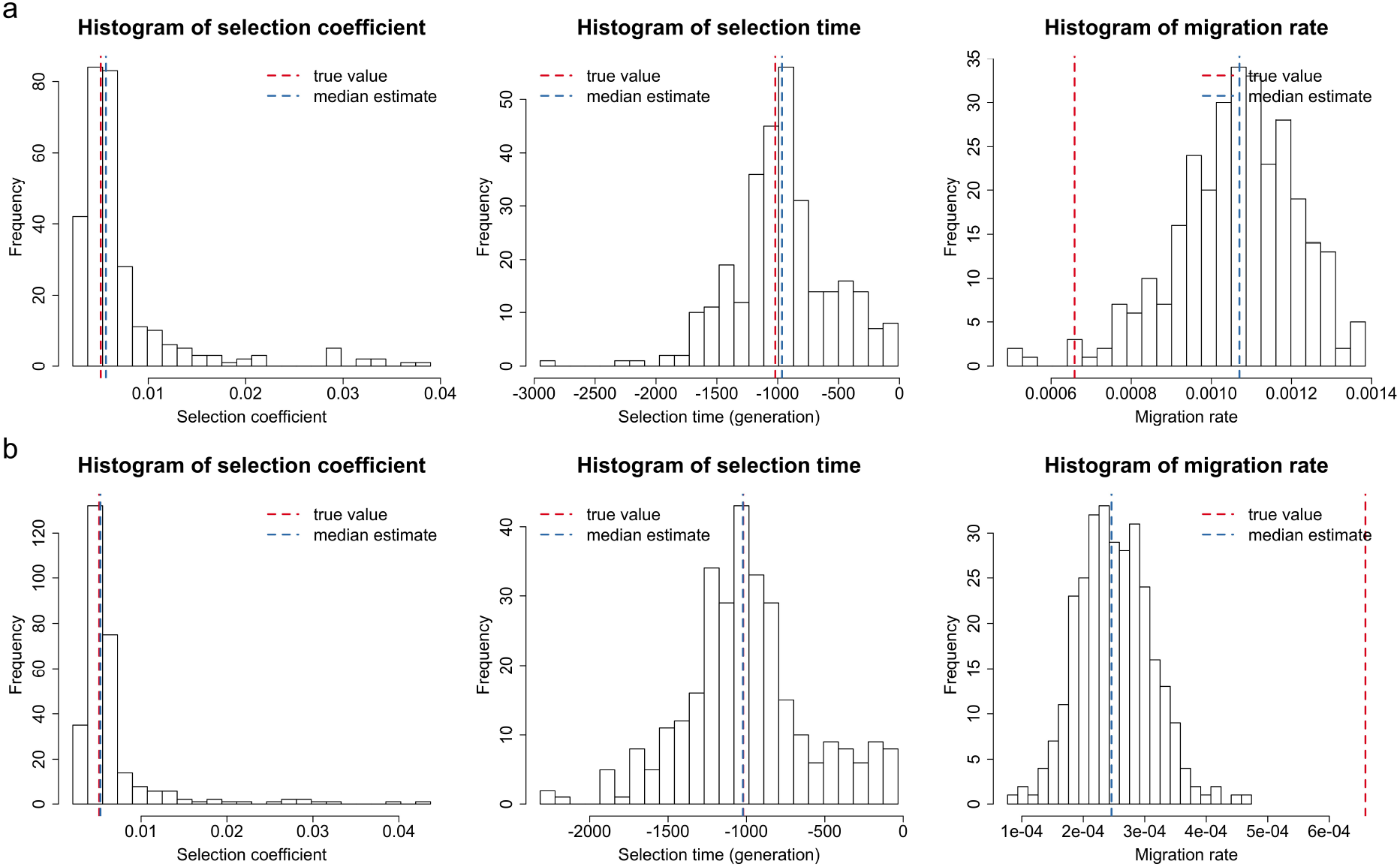
Empirical distributions of the estimates for 300 datasets simulated for *TSHR* based on the aDNA data presented in Table 2. We take the timing and strength of selection and migration to be those estimated with the population size *N* = 180000 given in Table 3, but the true migration time in the simulation is taken to be (a) *k*_*m*_ = −400 and (b) *k*_*m*_ = −100, respectively. To aid visual comparison, we have picked the *x* axis in the left panel not to cover all 300 estimates. The histogram containing all 300 estimates can be found in Figure S12.

In this work, we have focused on the continent-island model under the assumption that the allele frequencies of the continent population are fixed over time. As has been previously noted in the context of methods for detecting local adaptation (Lotterhos & Whitlock, 2015), caution must be exercised when applying to scenarios outside those that are validated in this study. Researchers may straightforwardly simulate test datasets under models that more closely reflect the assumptions of their study system (Haller & Messer, 2019) to investigate the robustness of our approach for their data.

Our Bayesian framework lends itself to being extended to more complex models of selection and migration. For example, we can allow the continent population to evolve under the Wright-Fisher diffusion with selection, therefore enabling us to model genetic drift and natural selection in the continent population. In this scenario, we need to simulate the underlying allele frequency trajectories of the continent population while we simulate those of the island population in our PMMH. If the continent population has been well studied, *i.e*., all required population genetic quantities can be pre-specified, our method is expected to have similar performance to this work. Otherwise, time serial samples from the continent population are required so that our method can be extended to the joint inference of selection acting on the continent population, where the likelihood will depend on the samples from both the island and continent populations, and the selection-related parameters for the continent population are updated as an additional block. In a similar manner, we can also allow gene migration to change the genetic composition of the continent population, *i.e*., the two-island model. Our approach is also readily applicable to the case of time-varying demographic histories such as Schraiber et al. (2016) and He et al. (2020c), but it may suffer from particle degeneracy and impoverishment issues if we extend our method to jointly estimate the allele age, which results from low-frequency mutant alleles at the early stage facing a higher probability of being lost.

It is possible to extend our procedure to handle the case of multiple islands or multiple loci. For multiple islands, our method will be more computationally demanding with an increase in the number of demes, but improvements to exact-approximate particle filtering techniques such as the PMMH algorithm continue to be developed (see, *e.g*., Yıldırım et al., 2018). For multiple (independent) loci, computational costs can be greatly reduced by updating the selection-related parameters for different loci on different cores in parallel. Our approach can be readily extended to the case of two linked loci by incorporating the method of He et al. (2020b), where modelling local linkage among loci has been illustrated to be capable of further improving the inference of selection, but such an extension will probably be computationally prohibitive in the case of multiple linked loci. As a tractable alternative for multiple linked loci, we can use our two-locus method in a pairwise manner by adding additional blocks in blockwise sampling.

## Acknowledgements

We are grateful to the communicating editors and the anonymous reviewers for their helpful comments on the earlier version of this work.

## Data Accessibility Statement

The authors state that all data necessary for confirming the conclusions of this work are represented fully within the article. Source code implementing the method described in this work is available at https://github.com/zhangyi-he/WFM-1L-DiffusApprox-PMMH-Chicken.

## Author Contributions

F.Y. and Z.H. designed the project and developed the method; W.L. and Z.H. implemented the method; W.L. and X.D. analysed the data under the supervision of M.B., F.Y. and Z.H.; W.L., X.D. and Z.H. wrote the manuscript; M.B. and F.Y. reviewed the manuscript.

