## Supplemental Information for "Inferring the timing and strength of natural selection and gene migration in the evolution of chicken from ancient DNA data"

#### File S1. Multi-allele Wright-Fisher model with selection and migration

Let $\boldsymbol{X}^{(N)}(k) = (X_1^{(N)}(k), X_2^{(N)}(k), X_3^{(N)}(k), X_4^{(N)}(k))$ denote the frequencies of the $\mathcal{A}_1^i$, $\mathcal{A}_2^i$, $\mathcal{A}_1^c$ and $\mathcal{A}_2^c$ alleles in $N$ zygotes of generation $k \in \mathbb{N}$ on the island. As discussed in Loog et al. (2017), the population dynamics might not be influenced by selection and migration in the early stage, thereby necessitating two additional population genetic quantities, the starting times of selection and migration on the island, denoted by $k_s$ and $k_m$, respectively. We then rewrite the selection coefficient and the migration rate as

$$s(k) = \begin{cases} 0, & \text{if } k < k_s \\ s, & \text{otherwise} \end{cases} \qquad \text{and} \qquad m(k) = \begin{cases} 0, & \text{if } k < k_m \\ m, & \text{otherwise,} \end{cases}$$

where the selection coefficient $s$ and the migration rate $m$ are both fixed from the timing of the onset of selection and migration up to present. Conditional on the allele frequencies of the population at generation $k$, we have

$$\boldsymbol{X}^{(N)}(k+1) \mid \boldsymbol{X}^{(N)}(k) = \boldsymbol{x} \sim \frac{1}{2N} \operatorname{Multinomial}(2N, \boldsymbol{p}),$$

where

$$\begin{aligned}
p_1 &= (1-m(k))x_1 \frac{1 - hs(k)(x_2+x_4)}{1 - 2hs(k)(x_1+x_3)(x_2+x_4) - s(k)(x_2+x_4)^2} \\
p_2 &= (1-m(k))x_2 \frac{1 - hs(k)(x_1+x_3) - s(k)(x_2+x_4)}{1 - 2hs(k)(x_1+x_3)(x_2+x_4) - s(k)(x_2+x_4)^2} \\
p_3 &= (1-m(k))x_3 \frac{1 - hs(k)(x_2+x_4)}{1 - 2hs(k)(x_1+x_3)(x_2+x_4) - s(k)(x_2+x_4)^2} + m(k)x_c \\
p_4 &= (1-m(k))x_4 \frac{1 - hs(k)(x_1+x_3) - s(k)(x_2+x_4)}{1 - 2hs(k)(x_1+x_3)(x_2+x_4) - s(k)(x_2+x_4)^2} + m(k)(1-x_c)
\end{aligned} \tag{1}$$

are the allele frequencies of an effectively infinite population on the island after random mating, selection and migration from generation $k$ to $k+1$. In Eq. (1), $x_c$ is the frequency of the $\mathcal{A}_1^c$ allele in the continent population, which is fixed over time.

We define the multi-allele Wright-Fisher model with selection and migration to be the random process $\boldsymbol{X}^{(N)} = \{\boldsymbol{X}^{(N)}(k), k \in \mathbb{N}\}$ evolving in the state space

$$\Omega_{\boldsymbol{X}^{(N)}} = \left\{ \boldsymbol{x} \in \left\{0, \frac{1}{2N}, \ldots, 1\right\}^4 : \sum_{i=1}^{4} x_i = 1 \right\}$$

with multinomial sampling probabilities described in Eq. (1). The transition probabilities of the allele frequencies from one generation to the next depend only on the current generation, which implies that the Wright-Fisher model $\boldsymbol{X}^{(N)}$ is a Markov process.

### File S2. Additional results for the analysis of simulated data

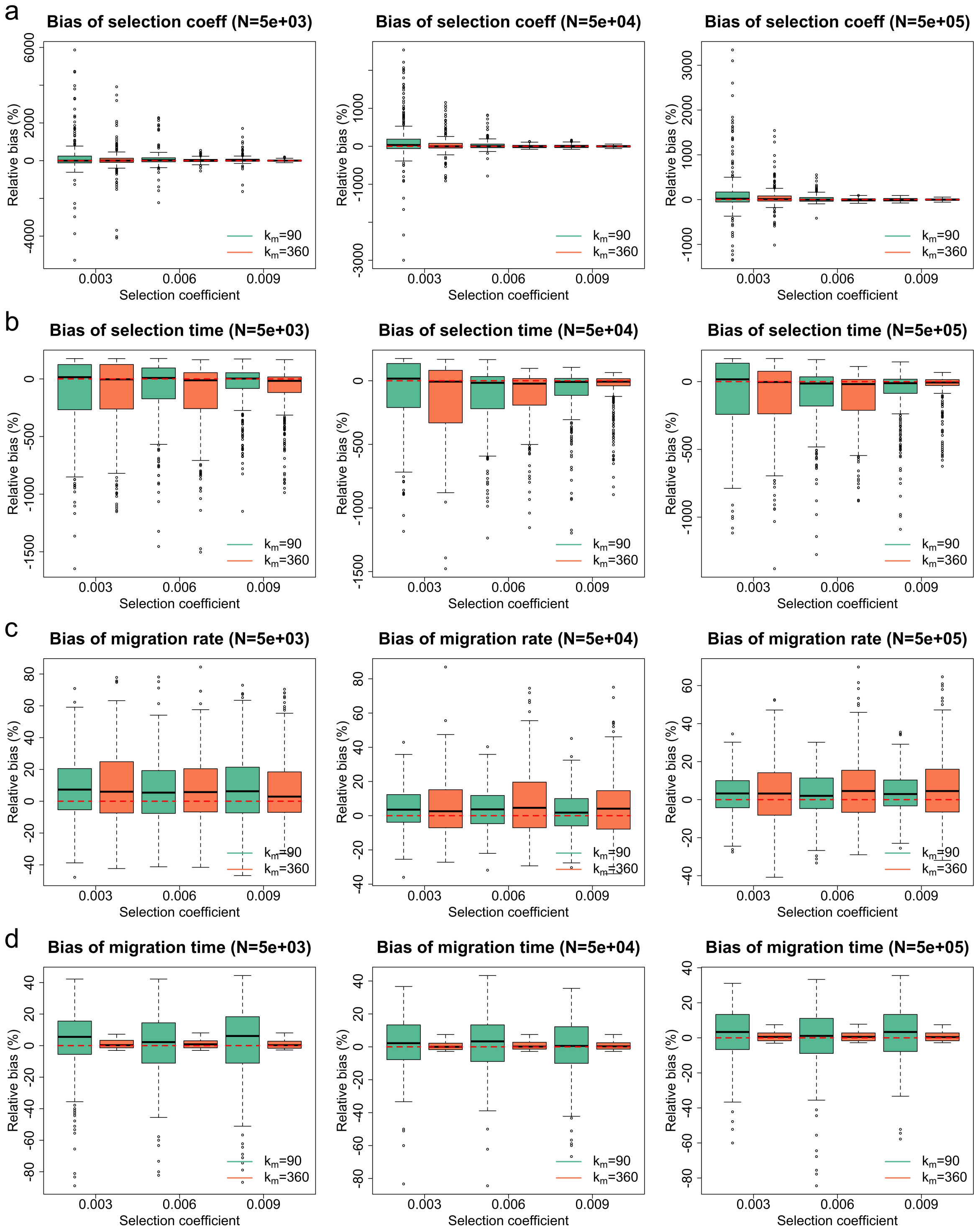


Figure S1: Empirical distributions of the estimates for 300 datasets simulated for additive selection ($h = 0.5$) where continent allele counts are not available at the first three sampling time points. Green boxplots represent the estimates produced for the case of selection starting after migration, and orange boxplots represent the estimates produced for the case of selection starting before migration. Boxplots of the relative bias of (a) the selection coefficient estimates (b) the selection time estimates (c) the migration rate estimates and (d) the migration time estimates.

| $N$ | $k_m$ | $s$ | Bias | Rel. Bias | RMSE | Rel. RMSE |
|---|---|---|---|---|---|---|
| 5000 | 90 | 0.003 | 0.00468 | 1.56050 | 0.02829 | 9.43106 |
| 5000 | 90 | 0.006 | 0.00502 | 0.83661 | 0.02561 | 4.26886 |
| 5000 | 90 | 0.009 | 0.00373 | 0.41479 | 0.02070 | 2.29946 |
| 5000 | 360 | 0.003 | 0.00093 | 0.31030 | 0.01953 | 6.51146 |
| 5000 | 360 | 0.006 | 0.00127 | 0.21204 | 0.00713 | 1.18760 |
| 5000 | 360 | 0.009 | 0.00045 | 0.05030 | 0.00491 | 0.54500 |
| 50000 | 90 | 0.003 | 0.00434 | 1.44527 | 0.01713 | 5.71086 |
| 50000 | 90 | 0.006 | 0.00157 | 0.26180 | 0.00805 | 1.34137 |
| 50000 | 90 | 0.009 | 0.00034 | 0.03819 | 0.00419 | 0.46556 |
| 50000 | 360 | 0.003 | 0.00100 | 0.33222 | 0.00682 | 2.27354 |
| 50000 | 360 | 0.006 | 0.00004 | 0.00686 | 0.00262 | 0.43607 |
| 50000 | 360 | 0.009 | −0.00008 | −0.00845 | 0.00260 | 0.28852 |
| 500000 | 90 | 0.003 | 0.00392 | 1.30796 | 0.01613 | 5.37626 |
| 500000 | 90 | 0.006 | 0.00099 | 0.16565 | 0.00540 | 0.89923 |
| 500000 | 90 | 0.009 | −0.00014 | −0.01584 | 0.00342 | 0.37963 |
| 500000 | 360 | 0.003 | 0.00156 | 0.52107 | 0.00700 | 2.33239 |
| 500000 | 360 | 0.006 | −0.00025 | −0.04204 | 0.00226 | 0.37589 |
| 500000 | 360 | 0.009 | −0.00005 | −0.00506 | 0.00225 | 0.24999 |

(a) Bias and RMSE of the selection coefficient estimates.

| $N$ | $k_m$ | $s$ | Bias | Rel. Bias | RMSE | Rel. RMSE |
|---|---|---|---|---|---|---|
| 5000 | 90 | 0.003 | −210.90333 | −1.17169 | 621.71914 | 3.45400 |
| 5000 | 90 | 0.006 | −156.32667 | −0.86848 | 521.88065 | 2.89934 |
| 5000 | 90 | 0.009 | −109.42000 | −0.60789 | 387.34442 | 2.15191 |
| 5000 | 360 | 0.003 | −211.54333 | −1.17524 | 597.03986 | 3.31689 |
| 5000 | 360 | 0.006 | −232.78333 | −1.29324 | 580.80568 | 3.22670 |
| 5000 | 360 | 0.009 | −181.31667 | −1.00731 | 430.67649 | 2.39265 |
| 50000 | 90 | 0.003 | −159.88000 | −0.88822 | 534.32684 | 2.96848 |
| 50000 | 90 | 0.006 | −204.42000 | −1.13567 | 478.61504 | 2.65897 |
| 50000 | 90 | 0.009 | −151.97333 | −0.84430 | 386.40670 | 2.14670 |
| 50000 | 360 | 0.003 | −221.46333 | −1.23035 | 555.05374 | 3.08363 |
| 50000 | 360 | 0.006 | −201.02000 | −1.11678 | 419.21672 | 2.32898 |
| 50000 | 360 | 0.009 | −114.76333 | −0.63757 | 308.31382 | 1.71285 |
| 500000 | 90 | 0.003 | −149.65667 | −0.83143 | 512.18792 | 2.84549 |
| 500000 | 90 | 0.006 | −178.32333 | −0.99069 | 429.41112 | 2.38562 |
| 500000 | 90 | 0.009 | −151.04333 | −0.83913 | 368.73452 | 2.04853 |
| 500000 | 360 | 0.003 | −189.66667 | −1.05370 | 509.13465 | 2.82853 |
| 500000 | 360 | 0.006 | −199.08333 | −1.10602 | 403.27948 | 2.24044 |
| 500000 | 360 | 0.009 | − 80.06333 | −0.44480 | 240.24256 | 1.33468 |

(b) Bias and RMSE of the selection time estimates.

Table S1: Bias and RMSE of the estimates for the selection-related parameters from 300 datasets simulated for additive selection ($h = 0.5$) where continent allele counts are not available at the first three sampling time points.

| $N$ | $k_m$ | $s$ | Bias | Rel. Bias | RMSE | Rel. RMSE |
|---|---|---|---|---|---|---|
| 5000 | 90 | 0.003 | 0.00040 | 0.07937 | 0.00105 | 0.21004 |
| 5000 | 90 | 0.006 | 0.00029 | 0.05825 | 0.00106 | 0.21144 |
| 5000 | 90 | 0.009 | 0.00036 | 0.07207 | 0.00115 | 0.22989 |
| 5000 | 360 | 0.003 | 0.00046 | 0.09101 | 0.00121 | 0.24261 |
| 5000 | 360 | 0.006 | 0.00041 | 0.08229 | 0.00114 | 0.22836 |
| 5000 | 360 | 0.009 | 0.00036 | 0.07261 | 0.00110 | 0.21985 |
| 50000 | 90 | 0.003 | 0.00020 | 0.03988 | 0.00062 | 0.12413 |
| 50000 | 90 | 0.006 | 0.00018 | 0.03647 | 0.00061 | 0.12148 |
| 50000 | 90 | 0.009 | 0.00012 | 0.02493 | 0.00061 | 0.12123 |
| 50000 | 360 | 0.003 | 0.00024 | 0.04795 | 0.00089 | 0.17748 |
| 50000 | 360 | 0.006 | 0.00036 | 0.07178 | 0.00101 | 0.20215 |
| 50000 | 360 | 0.009 | 0.00025 | 0.04997 | 0.00093 | 0.18545 |
| 500000 | 90 | 0.003 | 0.00015 | 0.03082 | 0.00056 | 0.11246 |
| 500000 | 90 | 0.006 | 0.00013 | 0.02547 | 0.00059 | 0.11724 |
| 500000 | 90 | 0.009 | 0.00019 | 0.03719 | 0.00057 | 0.11341 |
| 500000 | 360 | 0.003 | 0.00022 | 0.04329 | 0.00087 | 0.17372 |
| 500000 | 360 | 0.006 | 0.00030 | 0.05903 | 0.00088 | 0.17657 |
| 500000 | 360 | 0.009 | 0.00031 | 0.06182 | 0.00094 | 0.18709 |

(a) Bias and RMSE of the migration rate estimates.

| $N$ | $k_m$ | $s$ | Bias | Rel. Bias | RMSE | Rel. RMSE |
|---|---|---|---|---|---|---|
| 5000 | 90 | 0.003 | 2.41667 | 0.02685 | 19.15472 | 0.21283 |
| 5000 | 90 | 0.006 | 0.21333 | 0.00237 | 18.50171 | 0.20557 |
| 5000 | 90 | 0.009 | 1.71667 | 0.01907 | 21.02277 | 0.23359 |
| 5000 | 360 | 0.003 | 3.65333 | 0.01015 | 10.44956 | 0.02903 |
| 5000 | 360 | 0.006 | 3.70667 | 0.01030 | 10.05916 | 0.02794 |
| 5000 | 360 | 0.009 | 2.82333 | 0.00784 | 10.03278 | 0.02787 |
| 50000 | 90 | 0.003 | 1.77667 | 0.01974 | 14.92012 | 0.16578 |
| 50000 | 90 | 0.006 | 1.65333 | 0.01837 | 14.90011 | 0.16556 |
| 50000 | 90 | 0.009 | 0.14000 | 0.00156 | 15.82761 | 0.17586 |
| 50000 | 360 | 0.003 | 1.95667 | 0.00544 | 9.22551 | 0.02563 |
| 50000 | 360 | 0.006 | 2.50667 | 0.00696 | 9.32774 | 0.02591 |
| 50000 | 360 | 0.009 | 2.12667 | 0.00591 | 8.97552 | 0.02493 |
| 500000 | 90 | 0.003 | 1.62333 | 0.01804 | 14.32701 | 0.15919 |
| 500000 | 90 | 0.006 | −0.50333 | −0.00559 | 16.19311 | 0.17992 |
| 500000 | 90 | 0.009 | 2.09667 | 0.02330 | 13.71726 | 0.15241 |
| 500000 | 360 | 0.003 | 2.80333 | 0.00779 | 9.71820 | 0.02699 |
| 500000 | 360 | 0.006 | 2.71333 | 0.00754 | 9.71734 | 0.02699 |
| 500000 | 360 | 0.009 | 2.83333 | 0.00787 | 10.02630 | 0.02785 |

(b) Bias and RMSE of the migration time estimates.

Table S2: Bias and RMSE of the estimates for the migration-related parameters from 300 datasets simulated for additive selection ($h = 0.5$) where continent allele counts are not available at the first three sampling time points.

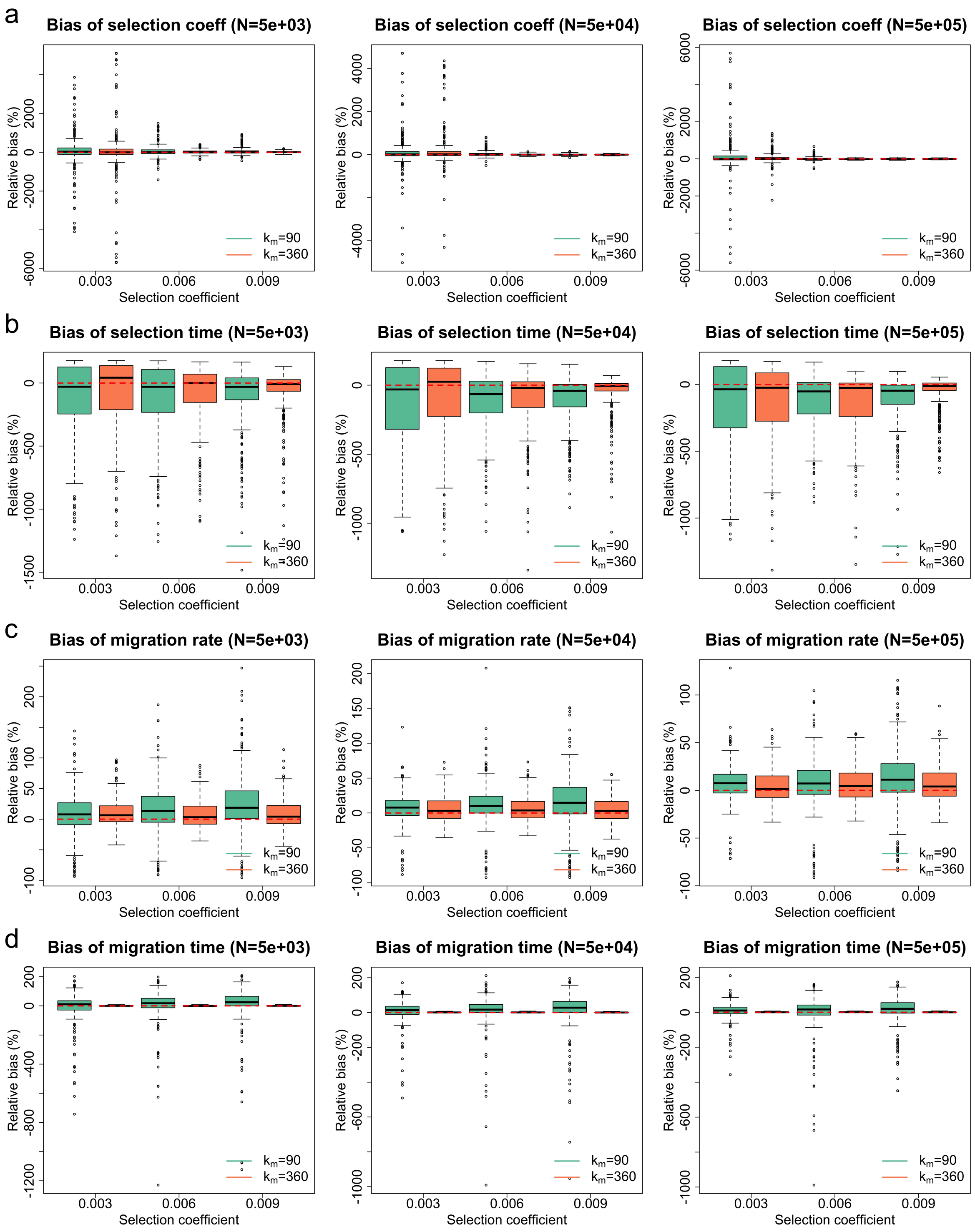


Figure S2: Empirical distributions of the estimates for 300 datasets simulated for additive selection ($h = 0.5$) where continent allele counts are not available at the first seven sampling time points. Green boxplots represent the estimates produced for the case of selection starting after migration, and orange boxplots represent the estimates produced for the case of selection starting before migration. Boxplots of the relative bias of (a) the selection coefficient estimates (b) the selection time estimates (c) the migration rate estimates and (d) the migration time estimates.

| $N$ | $k_m$ | $s$ | Bias | Rel. Bias | RMSE | Rel. RMSE |
|---:|---:|---:|---:|---:|---:|---:|
| 5000 | 90 | 0.003 | 0.00110 | 0.36598 | 0.02556 | 8.52108 |
| 5000 | 90 | 0.006 | 0.00347 | 0.57809 | 0.01812 | 3.02069 |
| 5000 | 90 | 0.009 | 0.00400 | 0.44399 | 0.01475 | 1.63932 |
| 5000 | 360 | 0.003 | 0.00046 | 0.15463 | 0.03540 | 11.80007 |
| 5000 | 360 | 0.006 | 0.00083 | 0.13876 | 0.00635 | 1.05828 |
| 5000 | 360 | 0.009 | 0.00093 | 0.10374 | 0.00472 | 0.52497 |
| 50000 | 90 | 0.003 | 0.00291 | 0.97150 | 0.02415 | 8.04869 |
| 50000 | 90 | 0.006 | 0.00171 | 0.28420 | 0.00768 | 1.27929 |
| 50000 | 90 | 0.009 | 0.00012 | 0.01355 | 0.00383 | 0.42527 |
| 50000 | 360 | 0.003 | 0.00524 | 1.74530 | 0.02562 | 8.54085 |
| 50000 | 360 | 0.006 | 0.00020 | 0.03408 | 0.00275 | 0.45771 |
| 50000 | 360 | 0.009 | −0.00009 | −0.01039 | 0.00250 | 0.27750 |
| 500000 | 90 | 0.003 | 0.00322 | 1.07406 | 0.03035 | 10.11574 |
| 500000 | 90 | 0.006 | 0.00088 | 0.14618 | 0.00622 | 1.03736 |
| 500000 | 90 | 0.009 | −0.00024 | −0.02651 | 0.00321 | 0.35614 |
| 500000 | 360 | 0.003 | 0.00144 | 0.48093 | 0.00901 | 3.00300 |
| 500000 | 360 | 0.006 | −0.00040 | −0.06742 | 0.00224 | 0.37318 |
| 500000 | 360 | 0.009 | −0.00049 | −0.05478 | 0.00237 | 0.26347 |

(a) Bias and RMSE of the selection coefficient estimates.

| $N$ | $k_m$ | $s$ | Bias | Rel. Bias | RMSE | Rel. RMSE |
|---:|---:|---:|---:|---:|---:|---:|
| 5000 | 90 | 0.003 | −218.83000 | −1.21572 | 613.45370 | 3.40808 |
| 5000 | 90 | 0.006 | −197.30000 | −1.09611 | 546.14176 | 3.03412 |
| 5000 | 90 | 0.009 | −179.02667 | −0.99459 | 483.29569 | 2.68498 |
| 5000 | 360 | 0.003 | −161.02000 | −0.89456 | 575.71969 | 3.19844 |
| 5000 | 360 | 0.006 | −147.86333 | −0.82146 | 459.69421 | 2.55386 |
| 5000 | 360 | 0.009 | −124.24000 | −0.69022 | 390.64411 | 2.17025 |
| 50000 | 90 | 0.003 | −235.89667 | −1.31054 | 586.78393 | 3.25991 |
| 50000 | 90 | 0.006 | −210.80000 | −1.17111 | 434.06003 | 2.41144 |
| 50000 | 90 | 0.009 | −188.10667 | −1.04504 | 366.53007 | 2.03628 |
| 50000 | 360 | 0.003 | −162.66333 | −0.90369 | 552.28193 | 3.06823 |
| 50000 | 360 | 0.006 | −182.22667 | −1.01237 | 419.28852 | 2.32938 |
| 50000 | 360 | 0.009 | − 98.90000 | −0.54944 | 284.07125 | 1.57817 |
| 500000 | 90 | 0.003 | −230.90000 | −1.28278 | 581.75415 | 3.23197 |
| 500000 | 90 | 0.006 | −218.24667 | −1.21248 | 420.77265 | 2.33763 |
| 500000 | 90 | 0.009 | −193.38333 | −1.07435 | 378.31113 | 2.10173 |
| 500000 | 360 | 0.003 | −214.52667 | −1.19181 | 533.35112 | 2.96306 |
| 500000 | 360 | 0.006 | −237.16333 | −1.31757 | 459.57492 | 2.55319 |
| 500000 | 360 | 0.009 | −108.98667 | −0.60548 | 256.16190 | 1.42312 |

(b) Bias and RMSE of the selection time estimates.

Table S3: Bias and RMSE of the estimates for the selection-related parameters from 300 datasets simulated for additive selection ($h = 0.5$) where continent allele counts are not available at the first seven sampling time points.

| $N$ | $k_m$ | $s$ | Bias | Rel. Bias | RMSE | Rel. RMSE |
|---|---|---|---|---|---|---|
| 5000 | 90 | 0.003 | 0.00046 | 0.09142 | 0.00182 | 0.36452 |
| 5000 | 90 | 0.006 | 0.00083 | 0.16636 | 0.00216 | 0.43224 |
| 5000 | 90 | 0.009 | 0.00132 | 0.26459 | 0.00275 | 0.54990 |
| 5000 | 360 | 0.003 | 0.00052 | 0.10371 | 0.00133 | 0.26544 |
| 5000 | 360 | 0.006 | 0.00038 | 0.07693 | 0.00116 | 0.23158 |
| 5000 | 360 | 0.009 | 0.00044 | 0.08795 | 0.00123 | 0.24646 |
| 50000 | 90 | 0.003 | 0.00035 | 0.06965 | 0.00118 | 0.23574 |
| 50000 | 90 | 0.006 | 0.00060 | 0.11954 | 0.00160 | 0.32043 |
| 50000 | 90 | 0.009 | 0.00084 | 0.16716 | 0.00203 | 0.40665 |
| 50000 | 360 | 0.003 | 0.00027 | 0.05318 | 0.00091 | 0.18284 |
| 50000 | 360 | 0.006 | 0.00032 | 0.06317 | 0.00098 | 0.19592 |
| 50000 | 360 | 0.009 | 0.00026 | 0.05257 | 0.00090 | 0.17914 |
| 500000 | 90 | 0.003 | 0.00035 | 0.07016 | 0.00107 | 0.21309 |
| 500000 | 90 | 0.006 | 0.00028 | 0.05678 | 0.00147 | 0.29372 |
| 500000 | 90 | 0.009 | 0.00064 | 0.12884 | 0.00177 | 0.35351 |
| 500000 | 360 | 0.003 | 0.00025 | 0.05038 | 0.00087 | 0.17486 |
| 500000 | 360 | 0.006 | 0.00033 | 0.06518 | 0.00095 | 0.18910 |
| 500000 | 360 | 0.009 | 0.00036 | 0.07105 | 0.00100 | 0.20017 |

(a) Bias and RMSE of the migration rate estimates.

| $N$ | $k_m$ | $s$ | Bias | Rel. Bias | RMSE | Rel. RMSE |
|---|---|---|---|---|---|---|
| 5000 | 90 | 0.003 | −6.86667 | −0.07630 | 96.89169 | 1.07657 |
| 5000 | 90 | 0.006 | 4.27667 | 0.04752 | 108.71174 | 1.20791 |
| 5000 | 90 | 0.009 | 7.58000 | 0.08422 | 136.62328 | 1.51804 |
| 5000 | 360 | 0.003 | 4.09667 | 0.01138 | 10.95491 | 0.03043 |
| 5000 | 360 | 0.006 | 2.77667 | 0.00771 | 9.73156 | 0.02703 |
| 5000 | 360 | 0.009 | 3.60333 | 0.01001 | 10.45227 | 0.02903 |
| 50000 | 90 | 0.003 | 4.36667 | 0.04852 | 60.81349 | 0.67571 |
| 50000 | 90 | 0.006 | 9.32333 | 0.10359 | 90.97170 | 1.01080 |
| 50000 | 90 | 0.009 | 11.73333 | 0.13037 | 108.29035 | 1.20323 |
| 50000 | 360 | 0.003 | 3.25667 | 0.00905 | 9.98916 | 0.02775 |
| 50000 | 360 | 0.006 | 3.64333 | 0.01012 | 10.28381 | 0.02857 |
| 50000 | 360 | 0.009 | 2.28333 | 0.00634 | 9.20344 | 0.02557 |
| 500000 | 90 | 0.003 | 6.21000 | 0.06900 | 45.52241 | 0.50580 |
| 500000 | 90 | 0.006 | −6.70333 | −0.07448 | 108.67571 | 1.20751 |
| 500000 | 90 | 0.009 | 13.19333 | 0.14659 | 78.06728 | 0.86741 |
| 500000 | 360 | 0.003 | 2.68333 | 0.00745 | 9.70550 | 0.02696 |
| 500000 | 360 | 0.006 | 3.45333 | 0.00959 | 10.37272 | 0.02881 |
| 500000 | 360 | 0.009 | 2.43333 | 0.00676 | 9.84412 | 0.02734 |

(b) Bias and RMSE of the migration time estimates.

Table S4: Bias and RMSE of the estimates for the migration-related parameters from 300 datasets simulated for additive selection ($h = 0.5$) where continent allele counts are not available at the first seven sampling time points.

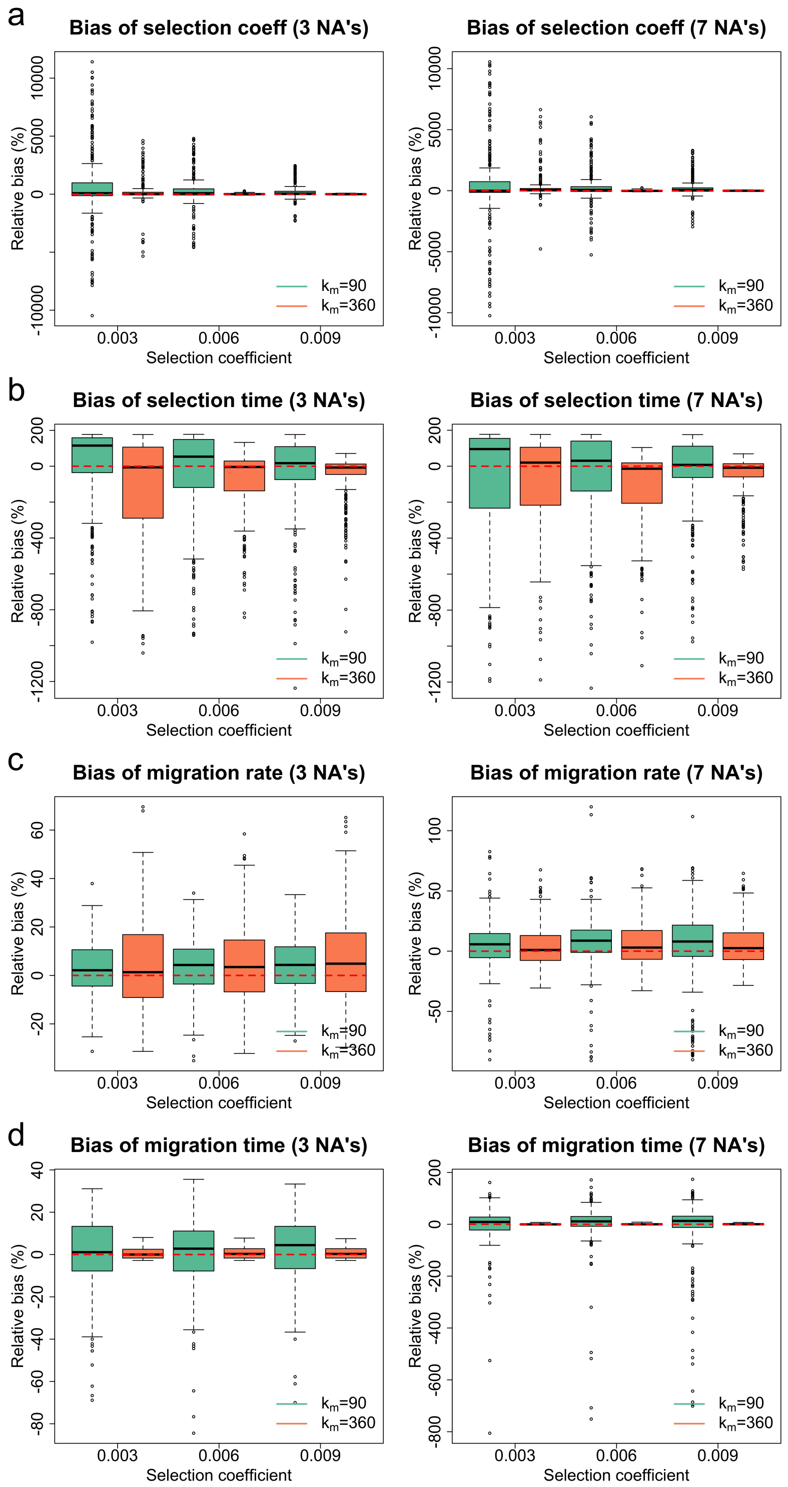


Figure S3: Empirical distributions of the estimates for 300 datasets simulated for dominant selection ($h = 0$) with the population size $N = 50000$. Green boxplots represent the estimates produced for the case of selection starting after migration, and orange boxplots represent the estimates produced for the case of selection starting before migration. Boxplots of the relative bias of (a) the selection coefficient estimates (b) the selection time estimates (c) the migration rate estimates and (d) the migration time estimates.

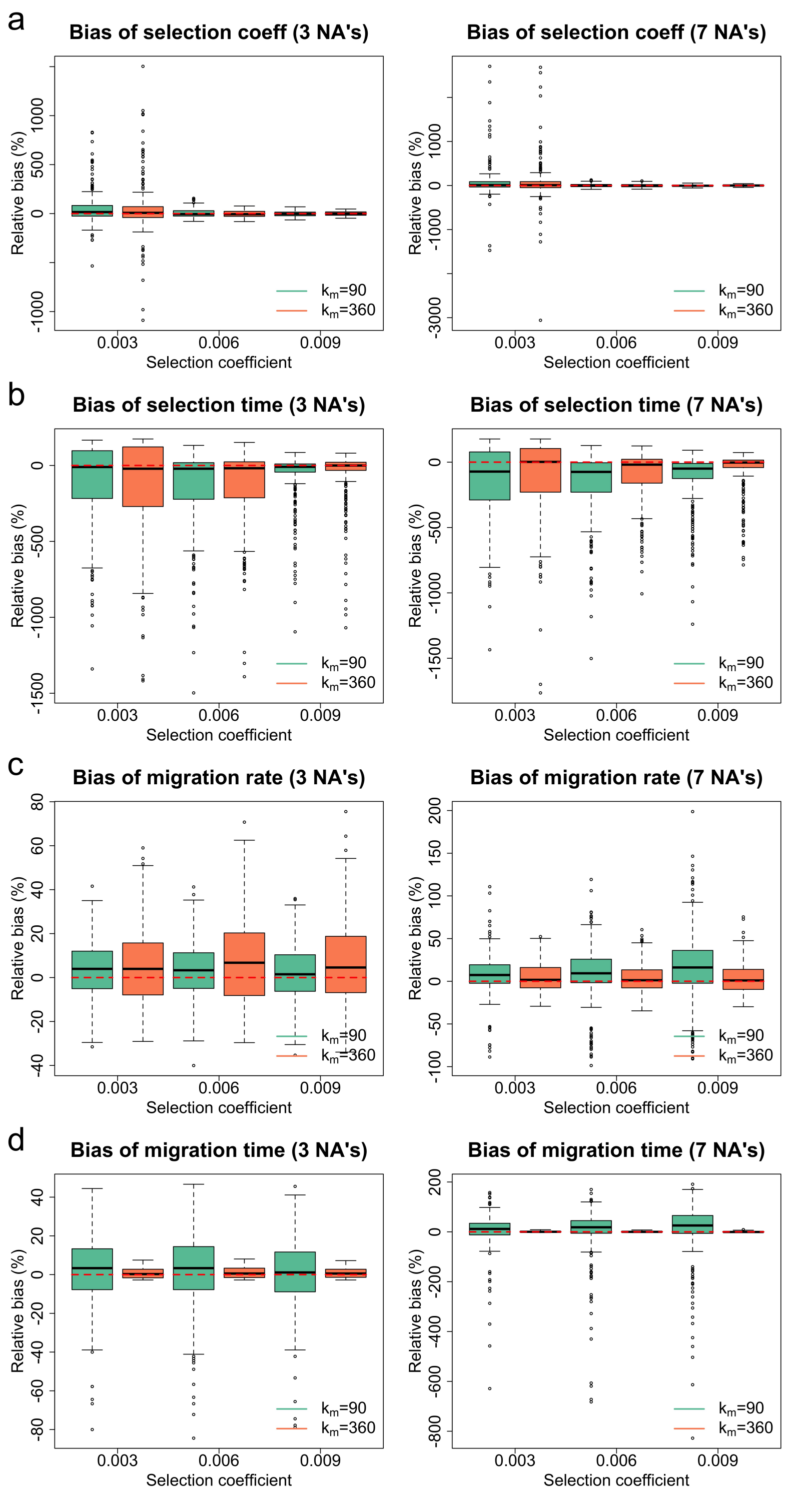


Figure S4: Empirical distributions of the estimates for 300 datasets simulated for recessive selection ($h = 1$) with the population size $N = 50000$. Green boxplots represent the estimates produced for the case of selection starting after migration, and orange boxplots represent the estimates produced for the case of selection starting before migration. Boxplots of the relative bias of (a) the selection coefficient estimates (b) the selection time estimates (c) the migration rate estimates and (d) the migration time estimates.

| Num. of NA's | $k_m$ | $s$ | Bias | Rel. Bias | RMSE | Rel. RMSE |
|---|---|---|---|---|---|---|
| 3 | 90 | 0.003 | 0.01973 | 6.57555 | 0.09331 | 31.10314 |
| 3 | 90 | 0.006 | 0.01975 | 3.29166 | 0.08571 | 14.28493 |
| 3 | 90 | 0.009 | 0.01877 | 2.08553 | 0.06014 | 6.68201 |
| 3 | 360 | 0.003 | 0.00500 | 1.66554 | 0.03001 | 10.00365 |
| 3 | 360 | 0.006 | 0.00109 | 0.18093 | 0.00413 | 0.68833 |
| 3 | 360 | 0.009 | 0.00002 | 0.00185 | 0.00338 | 0.37538 |
| 7 | 90 | 0.003 | 0.00969 | 3.23116 | 0.09536 | 31.78545 |
| 7 | 90 | 0.006 | 0.01830 | 3.04955 | 0.08103 | 13.50432 |
| 7 | 90 | 0.009 | 0.02142 | 2.38037 | 0.07270 | 8.07802 |
| 7 | 360 | 0.003 | 0.00864 | 2.87901 | 0.03119 | 10.39525 |
| 7 | 360 | 0.006 | 0.00008 | 0.01325 | 0.00331 | 0.55153 |
| 7 | 360 | 0.009 | −0.00044 | −0.04932 | 0.00289 | 0.32152 |

(a) Bias and RMSE of the selection coefficient estimates.

| Num. of NA's | $k_m$ | $s$ | Bias | Rel. Bias | RMSE | Rel. RMSE |
|---|---|---|---|---|---|---|
| 3 | 90 | 0.003 | 10.54667 | 0.05859 | 430.72468 | 2.39291 |
| 3 | 90 | 0.006 | − 72.46667 | −0.40259 | 467.49945 | 2.59722 |
| 3 | 90 | 0.009 | − 73.15000 | −0.40639 | 422.68432 | 2.34825 |
| 3 | 360 | 0.003 | −198.90333 | −1.10502 | 526.35467 | 2.92419 |
| 3 | 360 | 0.006 | −131.52667 | −0.73070 | 340.54389 | 1.89191 |
| 3 | 360 | 0.009 | −106.78667 | −0.59326 | 273.54705 | 1.51971 |
| 7 | 90 | 0.003 | −123.69667 | −0.68720 | 574.97869 | 3.19433 |
| 7 | 90 | 0.006 | − 92.04667 | −0.51137 | 480.65053 | 2.67028 |
| 7 | 90 | 0.009 | − 74.19333 | −0.41219 | 405.52244 | 2.25290 |
| 7 | 360 | 0.003 | −127.66000 | −0.70922 | 460.55611 | 2.55865 |
| 7 | 360 | 0.006 | −202.53333 | −1.12519 | 416.26197 | 2.31257 |
| 7 | 360 | 0.009 | −100.71000 | −0.55950 | 244.79350 | 1.35996 |

(b) Bias and RMSE of the selection time estimates.

Table S5: Bias and RMSE of the estimates for the selection-related parameters from 300 datasets simulated for dominant selection ($h = 0$) with the population size $N = 50000$.

| Num. of NA's | $k_m$ | $s$ | Bias | Rel. Bias | RMSE | Rel. RMSE |
|---|---|---|---|---|---|---|
| 3 | 90 | 0.003 | 0.00015 | 0.02925 | 0.00058 | 0.11688 |
| 3 | 90 | 0.006 | 0.00019 | 0.03765 | 0.00058 | 0.11594 |
| 3 | 90 | 0.009 | 0.00021 | 0.04117 | 0.00061 | 0.12270 |
| 3 | 360 | 0.003 | 0.00022 | 0.04440 | 0.00093 | 0.18662 |
| 3 | 360 | 0.006 | 0.00023 | 0.04590 | 0.00081 | 0.16230 |
| 3 | 360 | 0.009 | 0.00033 | 0.06627 | 0.00095 | 0.19083 |
| 7 | 90 | 0.003 | 0.00025 | 0.04953 | 0.00112 | 0.22334 |
| 7 | 90 | 0.006 | 0.00039 | 0.07708 | 0.00121 | 0.24102 |
| 7 | 90 | 0.009 | 0.00027 | 0.05307 | 0.00151 | 0.30259 |
| 7 | 360 | 0.003 | 0.00019 | 0.03860 | 0.00088 | 0.17604 |
| 7 | 360 | 0.006 | 0.00031 | 0.06215 | 0.00095 | 0.19017 |
| 7 | 360 | 0.009 | 0.00027 | 0.05400 | 0.00090 | 0.17993 |

(a) Bias and RMSE of the migration rate estimates.

| Num. of NA's | $k_m$ | $s$ | Bias | Rel. Bias | RMSE | Rel. RMSE |
|---|---|---|---|---|---|---|
| 3 | 90 | 0.003 | 0.48667 | 0.00541 | 15.83582 | 0.17595 |
| 3 | 90 | 0.006 | 0.94000 | 0.01044 | 15.46135 | 0.17179 |
| 3 | 90 | 0.009 | 1.73000 | 0.01922 | 14.71541 | 0.16350 |
| 3 | 360 | 0.003 | 2.14333 | 0.00595 | 9.88787 | 0.02747 |
| 3 | 360 | 0.006 | 2.92000 | 0.00811 | 9.93948 | 0.02761 |
| 3 | 360 | 0.009 | 2.78333 | 0.00773 | 10.27116 | 0.02853 |
| 7 | 90 | 0.003 | − 2.21333 | −0.02459 | 68.59538 | 0.76217 |
| 7 | 90 | 0.006 | 2.36000 | 0.02622 | 76.62780 | 0.85142 |
| 7 | 90 | 0.009 | −12.08667 | −0.13430 | 107.51425 | 1.19460 |
| 7 | 360 | 0.003 | 1.23667 | 0.00344 | 8.99981 | 0.02500 |
| 7 | 360 | 0.006 | 2.31333 | 0.00643 | 9.93177 | 0.02759 |
| 7 | 360 | 0.009 | 2.45333 | 0.00681 | 9.30340 | 0.02584 |

(b) Bias and RMSE of the migration time estimates.

Table S6: Bias and RMSE of the estimates for the migration-related parameters from 300 datasets simulated for dominant selection ($h = 0$) with the population size $N = 50000$.

| Num. of NA's | $k_m$ | $s$ | Bias | Rel. Bias | RMSE | Rel. RMSE |
|---|---|---|---|---|---|---|
| 3 | 90 | 0.003 | 0.00149 | 0.49529 | 0.00484 | 1.61276 |
| 3 | 90 | 0.006 | 0.00029 | 0.04762 | 0.00257 | 0.42811 |
| 3 | 90 | 0.009 | −0.00018 | −0.02048 | 0.00230 | 0.25598 |
| 3 | 360 | 0.003 | 0.00135 | 0.45071 | 0.00737 | 2.45705 |
| 3 | 360 | 0.006 | −0.00014 | −0.02306 | 0.00205 | 0.34194 |
| 3 | 360 | 0.009 | −0.00002 | −0.00178 | 0.00192 | 0.21367 |
| 7 | 90 | 0.003 | 0.00200 | 0.66651 | 0.01000 | 3.33475 |
| 7 | 90 | 0.006 | 0.00007 | 0.01223 | 0.00232 | 0.38643 |
| 7 | 90 | 0.009 | −0.00032 | −0.03524 | 0.00214 | 0.23779 |
| 7 | 360 | 0.003 | 0.00178 | 0.59398 | 0.01217 | 4.05581 |
| 7 | 360 | 0.006 | −0.00010 | −0.01692 | 0.00228 | 0.37986 |
| 7 | 360 | 0.009 | −0.00008 | −0.00913 | 0.00185 | 0.20605 |

(a) Bias and RMSE of the selection coefficient estimates.

| Num. of NA's | $k_m$ | $s$ | Bias | Rel. Bias | RMSE | Rel. RMSE |
|---|---|---|---|---|---|---|
| 3 | 90 | 0.003 | −180.96333 | −1.00535 | 510.54807 | 2.83638 |
| 3 | 90 | 0.006 | −240.48000 | −1.33600 | 511.44660 | 2.84137 |
| 3 | 90 | 0.009 | −112.56667 | −0.62537 | 312.30982 | 1.73505 |
| 3 | 360 | 0.003 | −225.53000 | −1.25294 | 596.65274 | 3.31474 |
| 3 | 360 | 0.006 | −223.48333 | −1.24157 | 483.37603 | 2.68542 |
| 3 | 360 | 0.009 | − 94.23333 | −0.52352 | 312.18813 | 1.73438 |
| 7 | 90 | 0.003 | −252.66333 | −1.40369 | 553.26892 | 3.07372 |
| 7 | 90 | 0.006 | −287.72333 | −1.59846 | 523.48826 | 2.90827 |
| 7 | 90 | 0.009 | −210.42333 | −1.16902 | 408.71711 | 2.27065 |
| 7 | 360 | 0.003 | −178.86667 | −0.99370 | 555.75419 | 3.08752 |
| 7 | 360 | 0.006 | −179.89333 | −0.99941 | 392.03727 | 2.17798 |
| 7 | 360 | 0.009 | −106.38000 | −0.59100 | 301.23072 | 1.67350 |

(b) Bias and RMSE of the selection time estimates.

Table S7: Bias and RMSE of the estimates for the selection-related parameters from 300 datasets simulated for recessive selection ($h = 1$) with the population size $N = 50000$.

| Num. of NA's | $k_m$ | $s$ | Bias | Rel. Bias | RMSE | Rel. RMSE |
|---|---|---|---|---|---|---|
| 3 | 90 | 0.003 | 0.00019 | 0.03834 | 0.00064 | 0.12782 |
| 3 | 90 | 0.006 | 0.00016 | 0.03129 | 0.00065 | 0.13011 |
| 3 | 90 | 0.009 | 0.00011 | 0.02125 | 0.00063 | 0.12681 |
| 3 | 360 | 0.003 | 0.00030 | 0.05904 | 0.00091 | 0.18134 |
| 3 | 360 | 0.006 | 0.00037 | 0.07449 | 0.00103 | 0.20542 |
| 3 | 360 | 0.009 | 0.00032 | 0.06390 | 0.00097 | 0.19369 |
| 7 | 90 | 0.003 | 0.00040 | 0.07995 | 0.00126 | 0.25223 |
| 7 | 90 | 0.006 | 0.00050 | 0.10071 | 0.00163 | 0.32588 |
| 7 | 90 | 0.009 | 0.00089 | 0.17809 | 0.00221 | 0.44209 |
| 7 | 360 | 0.003 | 0.00023 | 0.04628 | 0.00088 | 0.17529 |
| 7 | 360 | 0.006 | 0.00021 | 0.04253 | 0.00086 | 0.17192 |
| 7 | 360 | 0.009 | 0.00019 | 0.03777 | 0.00091 | 0.18105 |

(a) Bias and RMSE of the migration rate estimates.

| Num. of NA's | $k_m$ | $s$ | Bias | Rel. Bias | RMSE | Rel. RMSE |
|---|---|---|---|---|---|---|
| 3 | 90 | 0.003 | 1.73333 | 0.01926 | 15.82045 | 0.17578 |
| 3 | 90 | 0.006 | 1.23000 | 0.01367 | 17.06019 | 0.18956 |
| 3 | 90 | 0.009 | 0.34667 | 0.00385 | 15.67354 | 0.17415 |
| 3 | 360 | 0.003 | 2.74333 | 0.00762 | 9.71648 | 0.02699 |
| 3 | 360 | 0.006 | 3.67667 | 0.01021 | 10.51935 | 0.02922 |
| 3 | 360 | 0.009 | 3.23333 | 0.00898 | 9.67919 | 0.02689 |
| 7 | 90 | 0.003 | 4.12667 | 0.04585 | 64.94131 | 0.72157 |
| 7 | 90 | 0.006 | 3.07667 | 0.03419 | 94.74262 | 1.05270 |
| 7 | 90 | 0.009 | 10.45667 | 0.11619 | 100.60810 | 1.11787 |
| 7 | 360 | 0.003 | 2.59000 | 0.00719 | 9.83955 | 0.02733 |
| 7 | 360 | 0.006 | 1.95333 | 0.00543 | 9.10494 | 0.02529 |
| 7 | 360 | 0.009 | 0.93333 | 0.00259 | 8.89344 | 0.02470 |

(b) Bias and RMSE of the migration time estimates.

Table S8: Bias and RMSE of the estimates for the migration-related parameters from 300 datasets simulated for recessive selection ($h = 1$) with the population size $N = 50000$

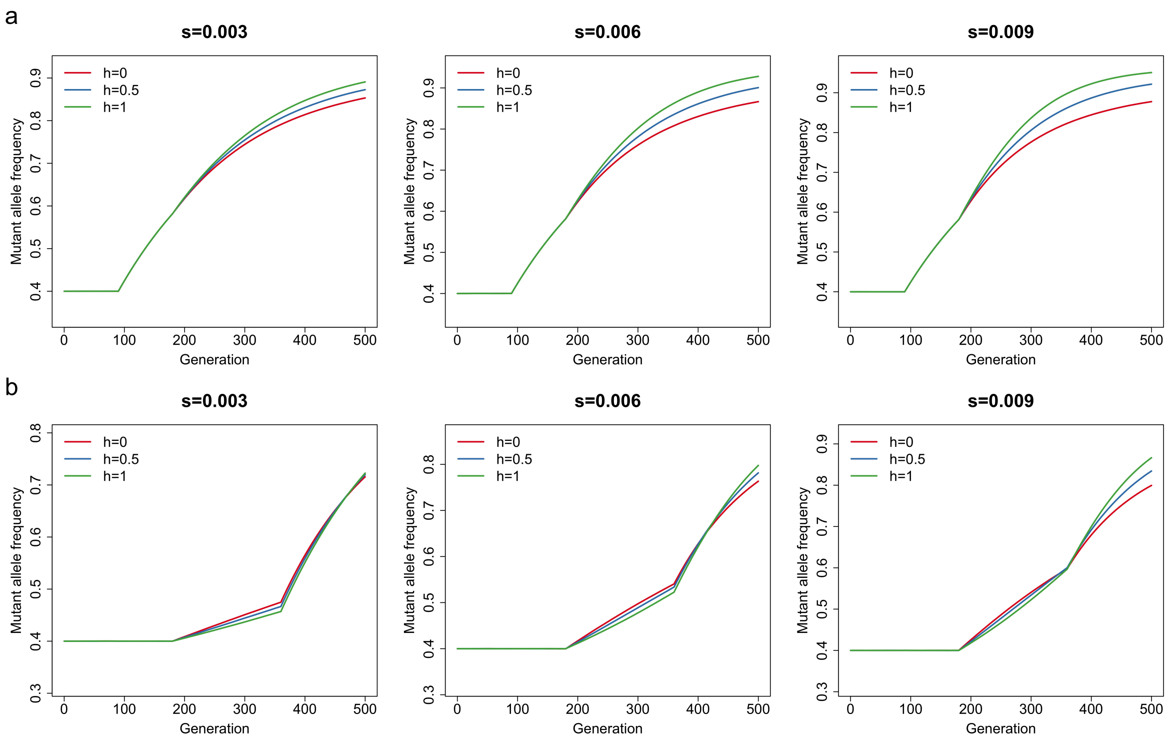


Figure S5: Mean mutant allele frequency trajectories of the underlying population for 18 possible combinations of the selection coefficient, the dominance parameter and the migration time. For each parameter combination, the mean trajectory of the mutant allele frequency is calculated with 10000 replicates. We take the population size to be $N = 50000$ and the migration time to be (a) $k_m = 90$ and (b) $k_m = 360$.

### File S3. Additional results for the analysis of real data

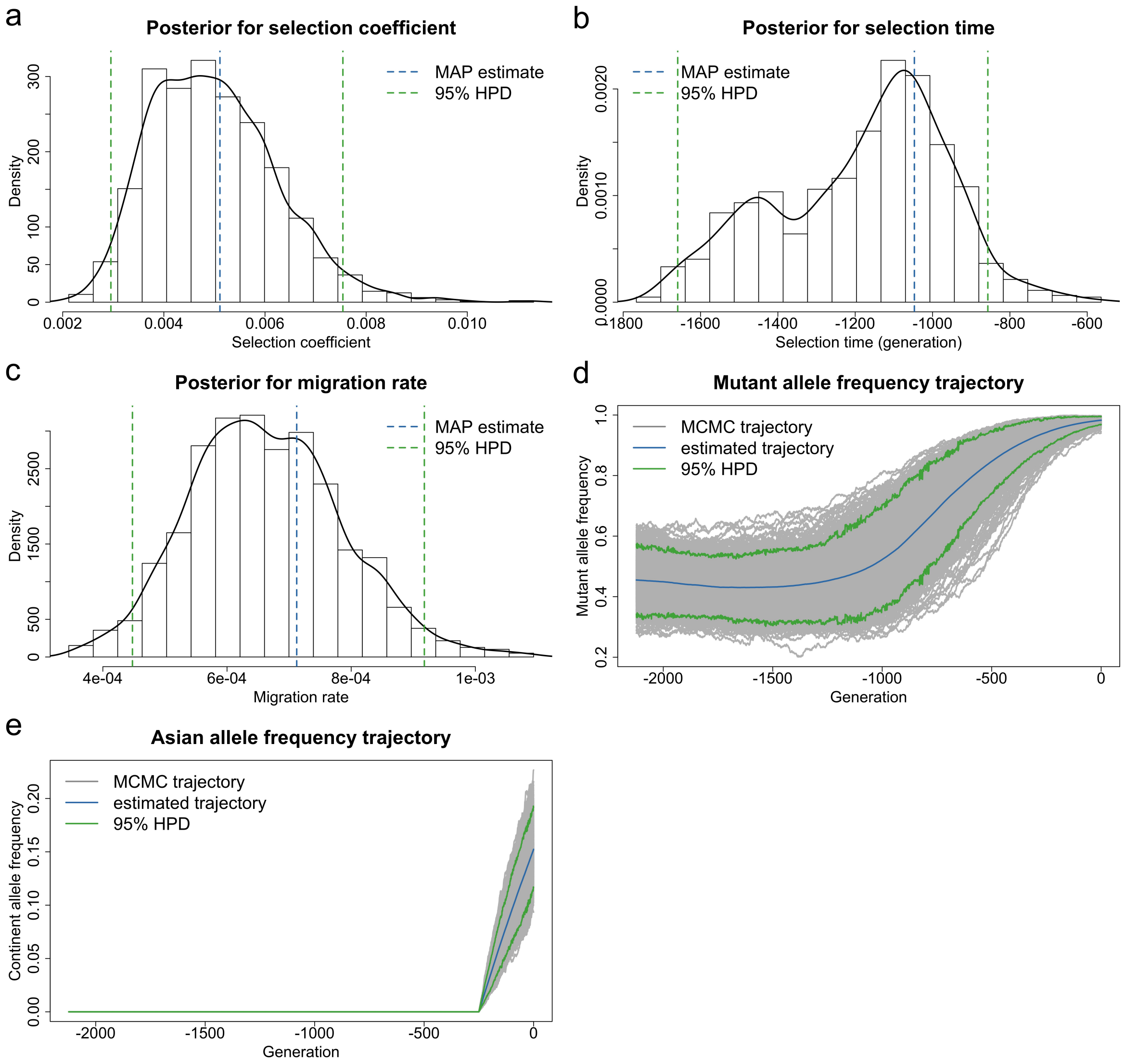


Figure S6: Bayesian estimates for aDNA data of European chicken genotyped at the *TSHR* locus from Loog et al. (2017) for the case of the population size $N = 26000$. Posteriors for (a) the selection coefficient (b) the selection time and (c) the migration rate. Estimated underlying trajectories of (d) the mutant allele frequency and (e) the Asian allele frequency in the European chicken population. The MAP estimate is for the joint posterior, and may not correspond to the mode of the marginals.

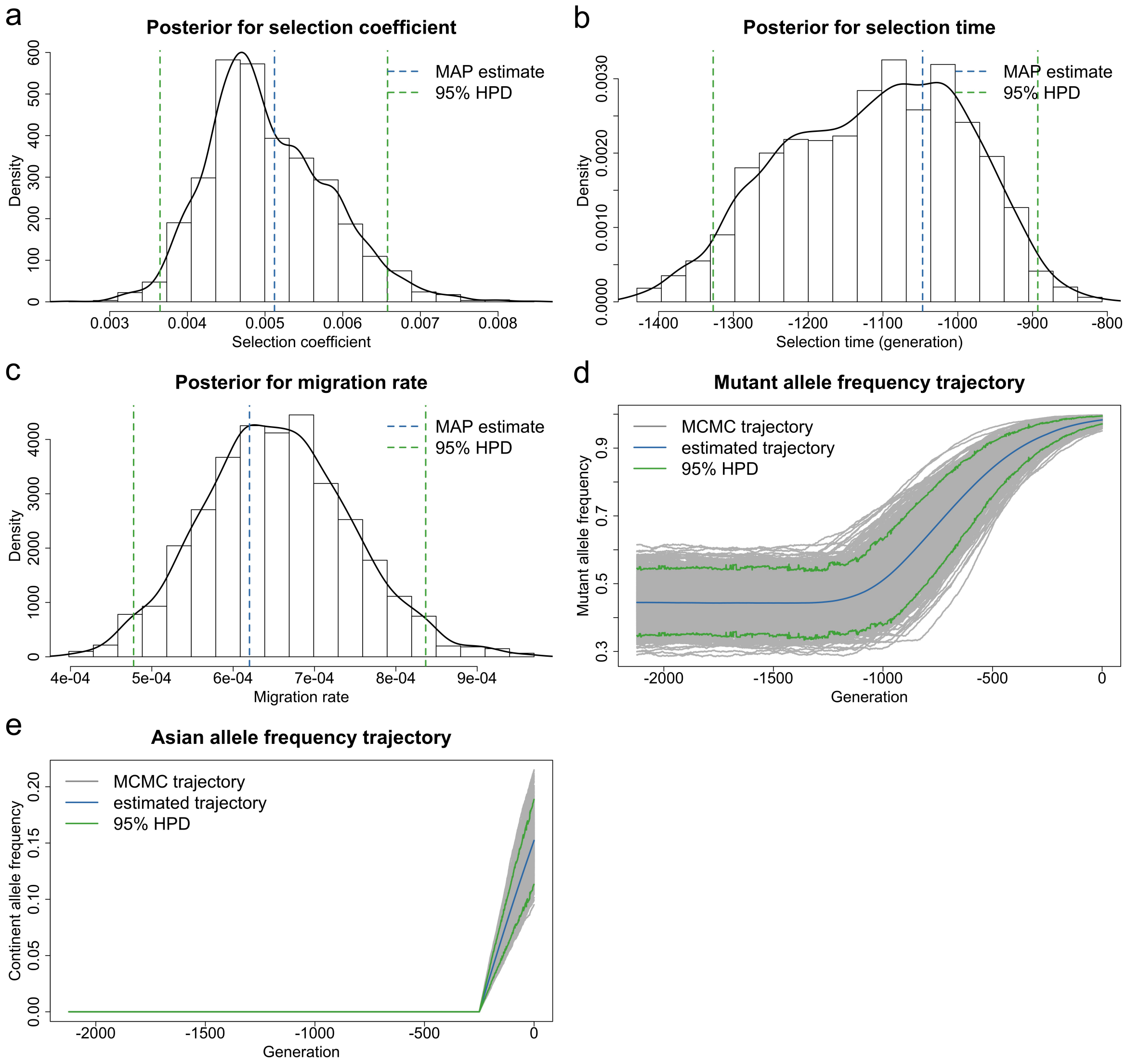


Figure S7: Bayesian estimates for aDNA data of European chicken genotyped at the *TSHR* locus from Loog et al. (2017) for the case of the population size $N = 460000$. Posteriors for (a) the selection coefficient (b) the selection time and (c) the migration rate. Estimated underlying trajectories of (d) the mutant allele frequency and (e) the Asian allele frequency in the European chicken population. The MAP estimate is for the joint posterior, and may not correspond to the mode of the marginals.

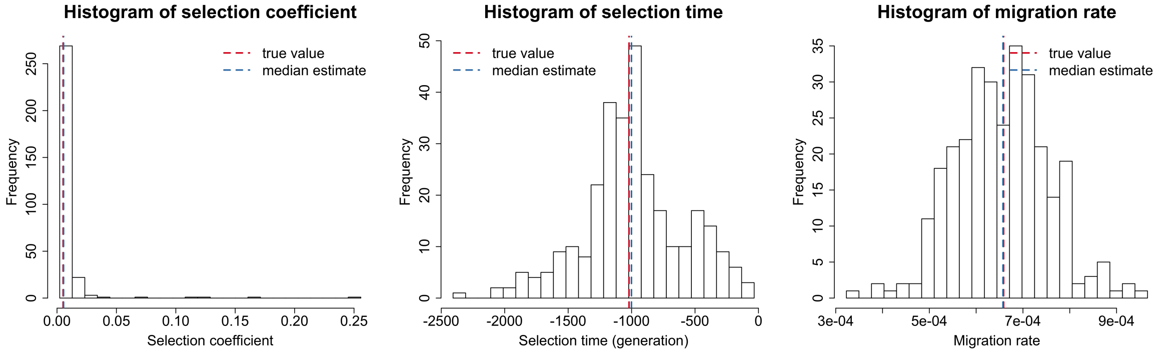


Figure S8: Empirical distributions of the estimates for 300 datasets simulated for *TSHR* based on the aDNA data shown in Table 2. We simulate the underlying population dynamics with the timing and strength of selection and migration estimated with the population size $N = 180000$ shown in Table 3.

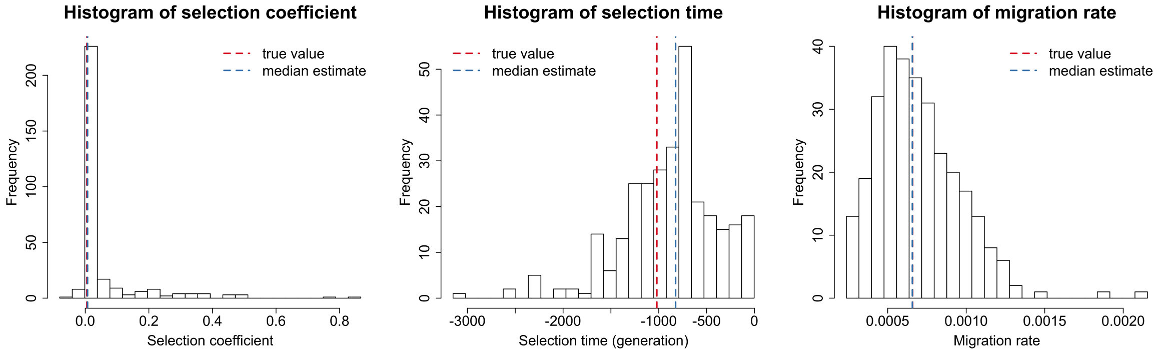


Figure S9: Empirical distributions of the estimates for 300 datasets simulated for *TSHR* based on the aDNA data presented in Table 2. We take the timing and strength of selection and migration to be those estimated with the population size $N = 180000$ given in Table 3, but the true population size in the simulation is taken to be $N = 4500$.

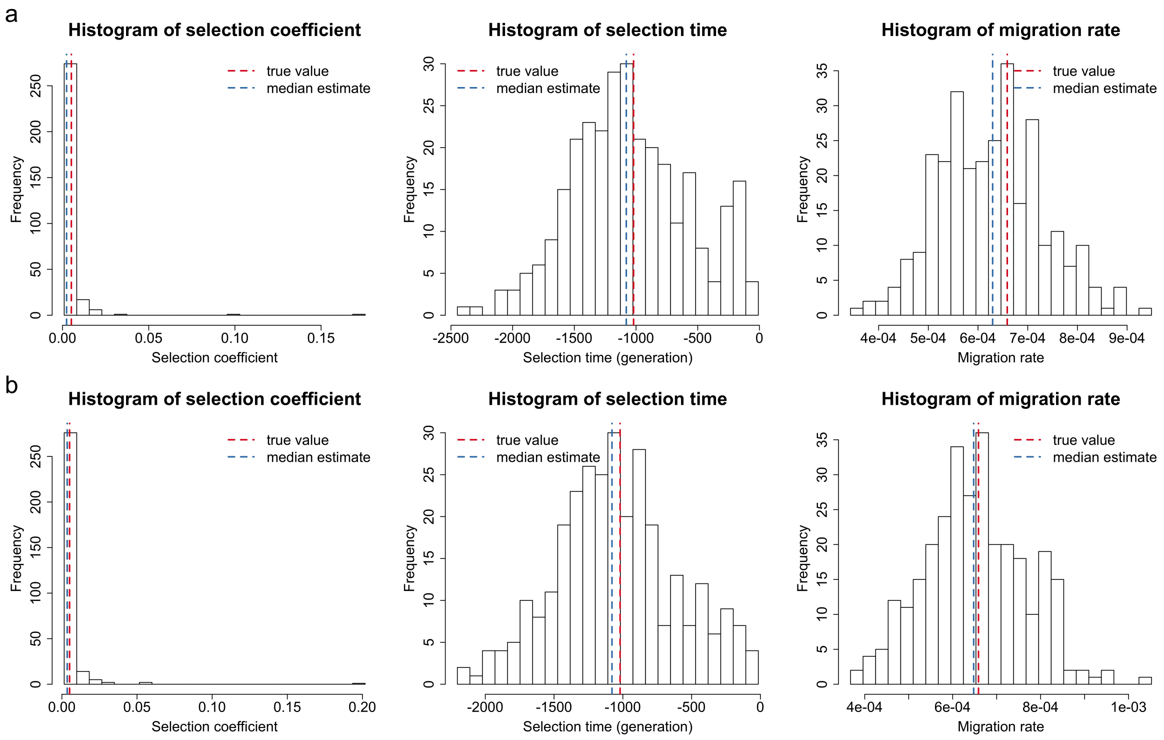


Figure S10: Empirical distributions of the estimates for 300 datasets simulated for *TSHR* based on the aDNA data presented in Table 2. We take the timing and strength of selection and migration to be those estimated with the population size $N = 180000$ given in Table 3, but the true dominance parameter in the simulation is taken to be (a) $h = 0$ and (b) $h = 0.5$, respectively.

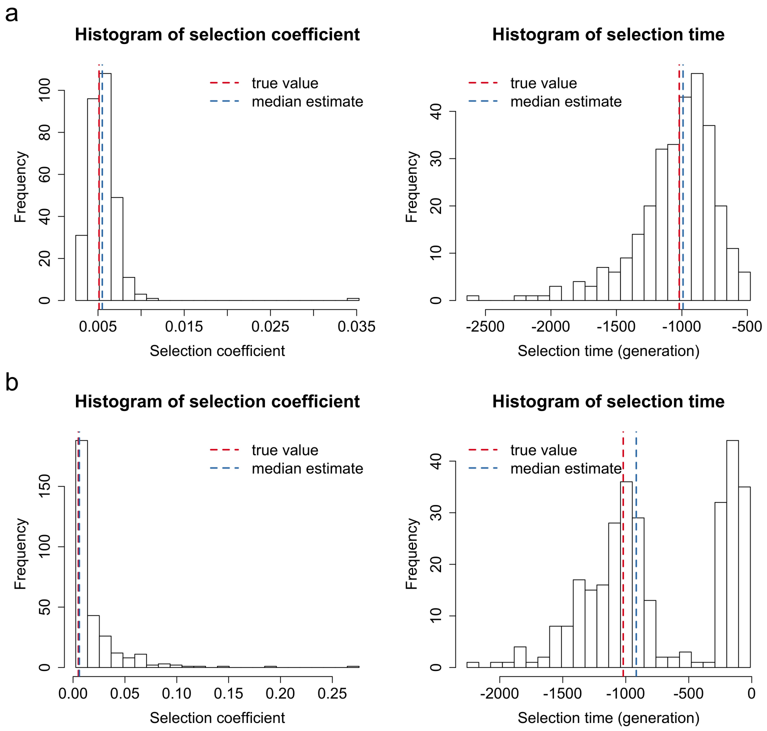


Figure S11: Empirical distributions of the estimates for 300 datasets simulated for *TSHR* based on the aDNA data presented in Table 2. We take the timing and strength of selection and migration to be those estimated with the population size $N = 180000$ given in Table 3, but the true migration rate in the simulation is taken to be (a) $m = 0.0001$ and (b) $m = 0.001$, respectively.

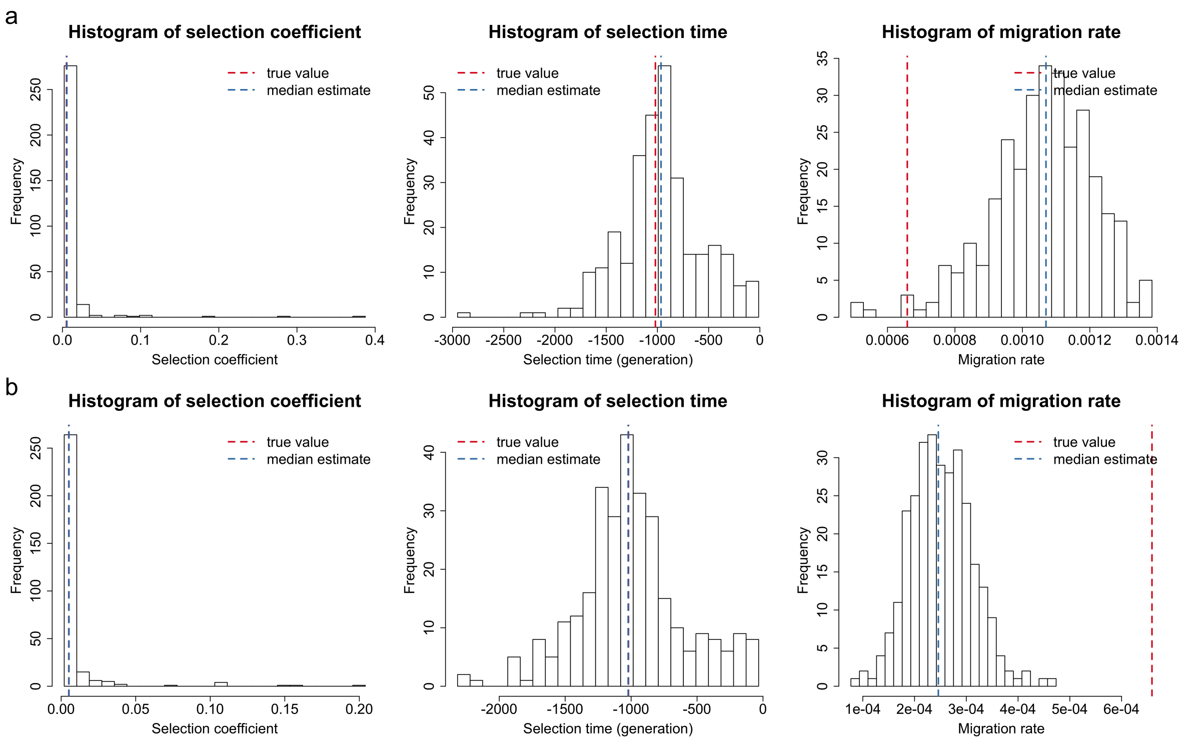


Figure S12: Empirical distributions of the estimates for 300 datasets simulated for *TSHR* based on the aDNA data presented in Table 2. We take the timing and strength of selection and migration to be those estimated with the population size $N = 180000$ given in Table 3, but the true migration time in the simulation is taken to be (a) $k_m = -400$ and (b) $k_m = -100$, respectively.
